# Integrative Bioinformatics Approach to Identify Prognostic Gene Signatures for Risk Stratification in Thyroid Carcinoma

**DOI:** 10.64898/2026.04.23.720344

**Authors:** Shivani Malik, Gajendra P. S. Raghava

## Abstract

Thyroid cancer is a heterogeneous malignancy with variable outcomes, highlighting the need for reliable biomarkers and effective risk stratification. In this study, we implemented a multi-step integrative framework to identify distinct prognostic biomarker sets using transcriptomic data from 572 thyroid cancer patients. Correlation analysis followed by false discovery rate (FDR) correction revealed significant associations of genes. Notably, MAFF (r = 0.25, p = 1.34×10⁻⁹, FDR = 2.46×10⁻⁷), NR4A3 (r = 0.24, p = 1.26×10⁻⁸, FDR = 9.25×10⁻⁷), and SRF showed strong positive correlations, whereas LOC728264 (r = –0.21, p = 7.39×10⁻⁷, FDR = 6.36×10⁻⁶) and VAMP1 (r = –0.20, p = 1.20×10⁻⁶, FDR = 1.3×10⁻⁴) exhibited negative correlations with OS. Univariate Cox regression identified several survival-associated genes, including TMEM90B (HR = 10.66, p = 2.88×10⁻⁵) and PTH1R (HR = 9.88, p = 5.55×10⁻⁵). LASSO regression further identified 31 key prognostic genes, including 13 potential drug targets predominantly functioning as inhibitors. Machine learning models based on seven independent 20-gene biomarker sets effectively predicted Class 0 (0–1 years), Class 1 (1–3 years), Class 2 (3–5 years), and Class 3 (>5 years), achieving AUC values of 0.91–0.94 and Kappa up to 0.76. An ensemble model further improved prediction (AUC = 0.95, Kappa = 0.72). Incorporating clinical variables (age, gender, stage) enhanced model performance (AUC = 0.96, Kappa = 0.80). Reduced 10- and 5-gene subsets demonstrated consistent yet slightly lower performance (AUC = 0.90 and 0.86, respectively). Collectively, the 20-gene set exhibited the strongest predictive and prognostic potential, highlighting the importance of integrating molecular and clinical features for risk stratification in thyroid cancer.All data and code are openly available (https://github.com/raghavagps/THCA_prognostic_biomarkers), supporting future research in thyroid cancer prediction.

## Introduction

Nowadays, the most prevalent endocrine cancer is thyroid carcinoma[1]. It is originating from thyroid parenchymal cells. While the mortality rate has remained relatively stable in recent years, the global incidence of thyroid cancer continues to rise. This disease presents a broad spectrum of clinical behaviors, ranging from slow-growing, indolent tumors to highly aggressive forms associated with significant mortality rates [2^i^]. According to GLOBOCAN estimates, there will be approximately 44,020 new cases of thyroid cancer and 2,290 related deaths in 2025 [3]. Thyroid carcinoma is more commonly diagnosed in women than men [4,5]. These statistics indicate the need for methods or biomarkers for detection of thyroid cancer. In this context, a reference technique is the thyroid nodule’s Fine Needle Aspiration (FNA) biopsy followed by cytological classification. FNA’s diagnostic accuracy has been found to be influenced by the operator’s expertise, the inherent features of nodules, and the interpretation of cytology [6,7]. One of the drawbacks is its poor capacity to detect follicular lesions. Owing to these FNA cytology limitations, a number of immunohistochemistry markers have been proposed; nevertheless, their effectiveness in diagnosing thyroid cancer is still being assessed.

The two main differentiated forms are the papillary (PTC) and follicular types (FTC). Medullary thyroid cancer (MTC) originates from calcitonin secreting C cells and is present in 3-5% of all thyroid cancers. Anaplastic thyroid cancer (ATC) is uncommon but highly aggressive [8]. Numerous investigations focused on possible factors causing increased TC incidence, such as chromosomal and genetic alterations, It seems that the activation of the mitogen-activated protein kinases (MAPK) and phosphoinositide 3 kinase-AKT (PI3K-AKT) signaling pathways might have a role in thyroid cancer growth [9] shown in **Figure 1**. In papillary thyroid cancer, activation of the MAPK signaling pathway occurs through two main mechanisms: recombination events and point mutations which are found in almost 70% of papillary cancers[10]. Chromosomal rearrangements such as rearrangement during transfection of proto-oncogene/papillary thyroid cancer (RET/PTC), PAX8/PPARγ, and B-type Raf kinase/A-kinase anchor protein 9 (BRAF/AKAP9) have been associated with exposure to ionizing radiation and likely represent fragile sites on the chromosome. Point mutations (RAS and BRAF genes) are most likely result of environmentally-induced or stochastic mutagenesis [10]. On the other hand, mutated RAS oncogene and PAX8/PPARγ fusion protein expression caused by translocation both enhance unregulated cell growth in 40% and 30% of subjects, respectively [11], and are expressed in follicular thyroid cancers. Other factors are iodine intake, TSH level, autoimmune thyroid disease, gender, estrogen, obesity, lifestyle changes, and environmental pollutants. Up to now, only childhood exposure to ionizing radiation has been fully recognized as a risk factor [12].

**Figure 1.**
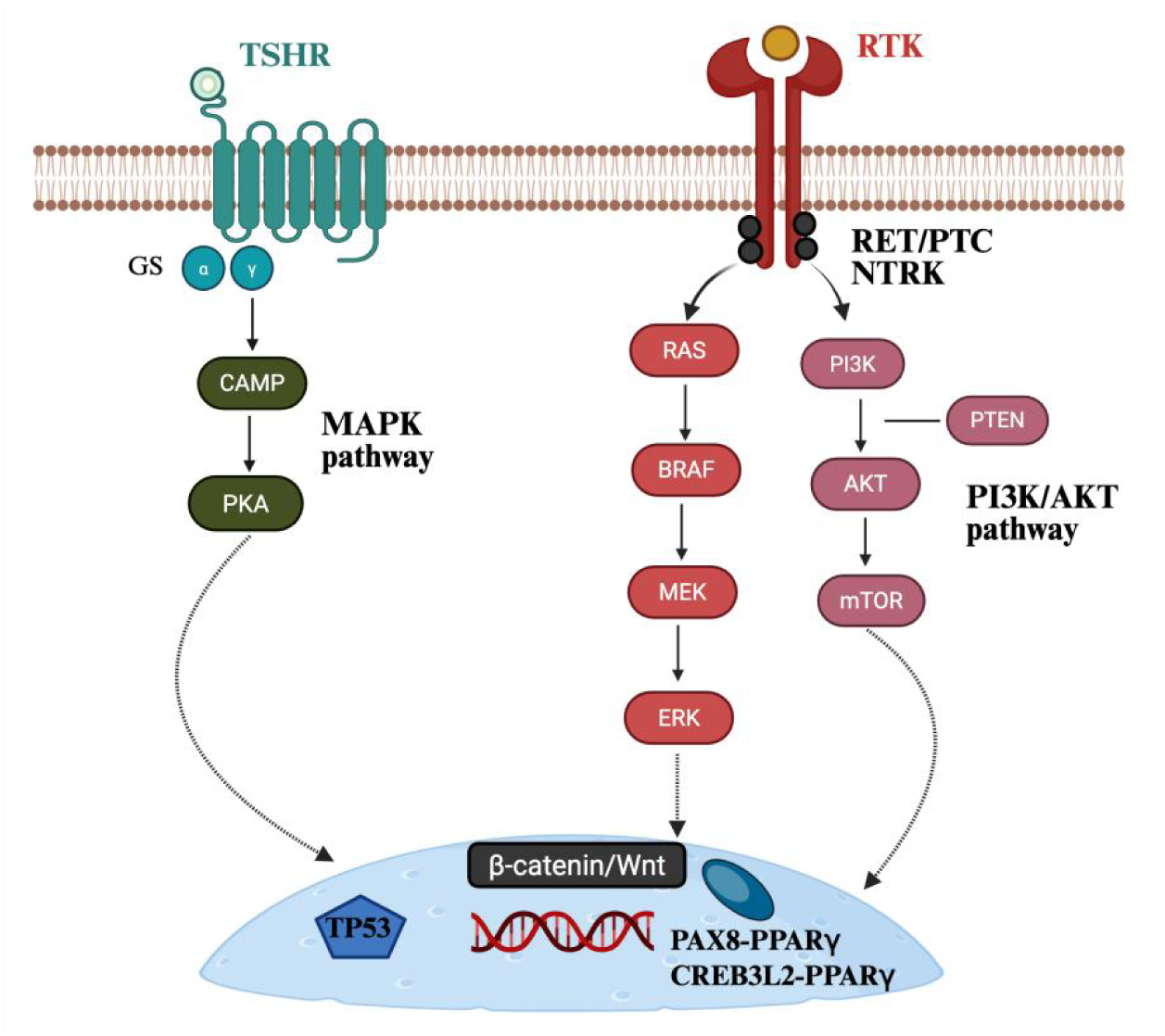
Key signaling pathways involved in thyroid cancer progression. TSHR activates the cAMP–PKA pathway via G-proteins, influencing transcriptional activity. The MAPK pathway is triggered by mutations in RAS, BRAF, or RET/PTC fusions, promoting cell proliferation through sequential activation of MEK and ERK. Additionally, the PI3K/AKT pathway, often altered through PI3K activation or PTEN loss, contributes to survival and growth via mTOR signaling. These cascades converge on nuclear targets such as TP53, β-catenin, and various transcription factors (e.g., PAX8–PPARγ), leading to tumor development.

Numerous studies focused on genes linked to pathways and on identification of diagnostic biomarkers. One of the study briefly revealed the nine key prognostic biomarker for predicting papillary thyroid carcinoma patients at high risk in apoptotic pathway with AUC of 0.92 [13]. In another study they found ADM, PXDN, MMP1, and TFF3 might serve as potential ATC diagnostic biomarkers with (AUC > 0.75) may be helpful for ATC molecular targeted therapy and immunotherapy[14]. In this study they identified a panel of 36 RNA transcripts that achieved F1 score of 0.75 with 0.73 AUROC (95% CI: 0.62–0.84) on the testing dataset. Moreover, prediction models based on 18-features from this panel correctly predicted 75% of the samples of the external validation dataset. In addition, the multiclass model classified normal, early, and late-stage samples with AUROC of 0.95 (95% CI: 0.84–1), 0.76 (95% CI: 0.66–0.85) and 0.72 (95% CI: 0.61–0.83)on the validation dataset[15]. Furthermore one of the study identified RXRG, CDH2, ETV5, QPCT, LRP4, FN1, and LPAR5” as PTC biomarkers, providing novel reference markers for the diagnosis and treatment of PTC patients[16]. In this study machine learning suggested potential diagnostic biomarkers for PDLIM4, ANXA1, PKM, NPC2, LMNA, and FN1. Finally, the five-protein nomogram may be a prognostic biomarker in the future[17]. In this study, we used LASSO-Cox regression to select six IRFGs, which were incorporated into the prognostic risk model. For machine learning analysis on the basis of risk score divide the patienst into high or low risk group they achieved AUCs for 1-year, 3-year, and 5-year OS were 0.971, 0.740, and 0.756 in training set and 0.621, 0.763, and 0.758 on validation set for risk stratification[18].

The motivation for integrating Machine Learning (ML) in thyroid cancer research stems from the limitations of conventional diagnostic and monitoring approaches, as ML offers transformative potential for reducing human errors and improving prediction outcomes for diagnostic accuracy, risk stratification, treatment options, recurrence prognosis, and patient quality of life[19]. Due to advent of high-throughput sequencing methods and public databases, many biomarkers have been identified for TC diagnosis, classification, and prognosis. In this study, we exploited the mRNA expression data obtained from The Cancer Genome Atlas-Thyroid Carcinoma (TCGA-THCA) cohort and identified distinct set of biomarkers that are associated with PTC prognosis.

As far as we are aware, these methods often only find one set of biomarkers. Furthermore, given a limited group of data, they usually select prognostic indicators, such as genes associated with the apoptotic process and the immunological response. Our work is a unique approach in which we attempt to construct biomarker sets from all genes, rather than focusing on a subset. Additionally, among the remaining genes, we select a secondary collection of biomarkers after identifying an initial set. By repeating this procedure, seven distinct sets of prognostic biomarkers are identified. Our study’s methodology is shown in **Figure 2**.

**Figure 2.**
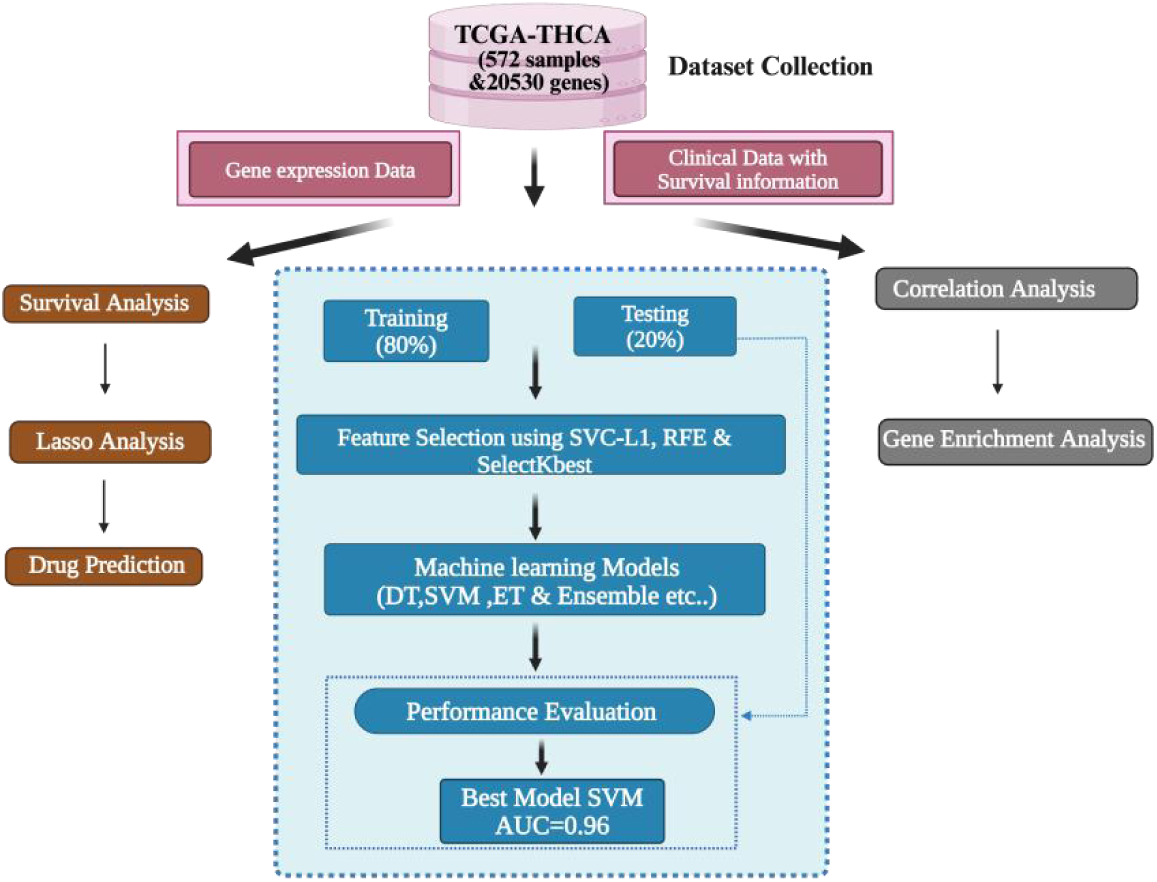
The workflow of our study

## Methodology

### Dataset Acquisition and Prepr ocessing

UCSC Xena (https://xenabrowser.net/datapages/) was used to acquire data from TCGA-THCA. Transcriptomic data comprising 20,530 genes is available for 572 samples, while survival information is available for 580 samples. Using a Python script, the preprocessing method first eliminated duplicate features, filtered out genes with more than 50% zero values, then eliminated low-variance features to ensure that only informative genes remained. We kept 10876 genes for further analysis after preprocessing. We found a cohort of 572 patients with available survival information after combining this gene expression data with the relevant clinical data. Then we scaled the data using a standard scaler[20]

### Statistical Analysis

#### Correlation Analysis

Pearson correlation analysis was performed to identify genes significantly associated with overall survival (OS) by comparing gene expression levels with survival time [21]. This was implemented in R using a custom loop that applied the cor() and cor.test() functions to compute correlation coefficients and p-values for each gene. To account for multiple testing, p-values were adjusted using the false discovery rate (FDR) method, and genes with FDR-adjusted p-values < 0.05 were considered significant. Genes showing significant positive correlations were selected for further analysis. To explore their potential functional roles in thyroid cancer, these genes were subjected to gene enrichment analysis [22]. Gene Ontology (GO) and Reactome pathway enrichment analyses were performed using the gseapy package in Python, with a significance cutoff of 0.05.

#### Survival Analysis

To assess the predictive importance of gene expression levels, survival analysis was conducted. Vital status (coded as 1 = dead, 0 = alive) and overall survival time were employed as outcome variables. For every gene, survival between groups with high and low expression was compared using Kaplan–Meier survival curves using survival and survwiner package in R with the median expression value serving as the cutoff. The statistical significance was evaluated using the log-rank test[23].

The coxph() function from the survival package in R was also used to estimate hazard ratios (HR) and 95% CI for each gene through univariate Cox proportional hazards regression. P-values < 0.05 were used to determine significant genes. Then, using the R glmnet package, LASSO regression analysis was carried out to find predictive genes, with an ideal lambda value of 0.03[24]. Depending on the formula:

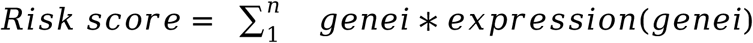

Then, using the Drug Gene Interaction Database (DGIdb) to suggest possible treatment agents based on known drug-gene interactions, identified prognostic genes were examined[25]

#### Machine Learning Analysis

For the prognostic model, we used a variety of feature selection strategies to find informative genes. Four groups of people were identified based on their overall survival time: Class 0 (0–1 years), Class 1 (1–3 years), Class 2 (3–5 years), and Class 3 (>5 years).

#### Feature Selection methods

We used a number of well-known feature selection techniques, such as Recursive Feature Elimination (RFE), Linear Support Vector Classification with L1 penalty (SVC-L1), and SelectKBest, to determine which characteristics were most important for the prognostic model. Similar investigations have effectively used these techniques[26,27,28].

#### Selection of Key Features for the Primary Biomarker Set

Using the Scikit-learn package, we used a number of feature selection techniques to determine the primary set of important biomarkers. Initially we employed a sparsity-inducing method called Linear Support Vector Classification (SVC) with L1 regularization, which gives less relevant features zero weights. To ensure a targeted and pertinent feature subset, we determined the absolute values of the SVC-L1 coefficients and kept features with scores higher than the average coefficient value. we also employed Recursive Feature Elimination (RFE), which picks an ideal subset by recursively eliminating the least significant features after ranking features according to relevance[29]. To extract the top-ranked features based on their statistical significance, we then used the SelectKBest technique using the f_classif scoring function.

#### Uncovering Complementary Sets of Biomarkers

To determine the secondary biomarker set, all features that were a part of the main set were eliminated before reapplying the SVC-L1 algorithm. Features of the recently selected biomarker set are independent and non-overlapping. Until we discovered seven sets of twenty prognostic biomarkers, this process was repeated.

#### Machine Learning Models

Using the Synthetic Minority Over-sampling Technique (SMOTE), the problem of class imbalance was addressed[30]. Stratified five-fold cross-validation was used to evaluate the model’s resilience and performance, with training and testing datasets split 80:20. A wide range of machine learning (ML) algorithms were used, such as k-Nearest Neighbors (KNN), Light Gradient Boosting Machine (LightGBM), Multi-Layer Perceptron (MLP), Decision Tree (DT), Extreme Gradient Boosting (XGB), Support Vector Machine (SVM), Extra Trees Classifier (ET), Random Forest (RF), and Logistic Regression (LR). Each of these methods was used to create prediction models across different feature sets in order to assess how well they performed in classification tasks[31].

#### Ensemble Models

We used both voting and stacking classifiers in our ensemble learning algorithms to improve prediction performance. Using a hard voting technique, we aggregated predictions from several basic classifiers (RF,ET, LightGBM, and SVM) in the voting ensemble. The final class was decided by the majority vote. By using this method, we were able to combine the advantages of various models to enhance generalization. In order to employ predictions from multiple base learners (such as RF, ET, and LGBM) as input features for a meta-classifier, we also constructed a stacking ensemble. First, we learned the best combination of base model predictions using SVM as the meta-model. The meta-model was able to learn higher-order patterns because to its two-tiered model architecture, which also improved overall prediction accuracy by successfully capturing relationships between the outputs of the base models.

#### Feature Selection for Reduced Biomarker Subsets

From each of the seven biomarker sets obtained, we further selected the top-ranked features based on their weights assigned by the SVC-L1 algorithm. Specifically, we extracted the top 10 and top 5 features from every set to evaluate whether smaller subsets of biomarkers could provide better or more efficient prognostic performance.

#### Integration of Clinical Features

To enhance the predictive capability of the models, clinical variables such as patient age, gender, and cancer stage were incorporated alongside gene expression features. Gender was encoded numerically (male = 1, female = 0), and cancer stage was converted into ordinal values (Stage I = 0, Stage II = 1, Stage III = 2, Stage IV = 3). Age was treated as a continuous variable and standardized using a standard scaler. These clinical features were combined with gene set to create extended feature sets, which were then used as input for machine learning model development. All preprocessing and integration steps were carried out in Python and R.

#### Prognostic Modeling of 20 and 10 Gene Sets Using Survival Analysis

To evaluate the prognostic significance of seven predefined 20-gene sets, we performed multivariate Cox proportional hazards modeling using gene expression data, overall survival time, and vital status. For each gene set, a risk score was computed as a linear combination of gene expressions weighted by Cox model coefficients. This risk score was then used in a univariate Cox model to obtain a single HR representing the prognostic impact of the gene set. To further refine the models, we also repeated this procedure using the top 10 and top 5 genes from each set. In addition, we examined the contribution of clinical factors by extending the models to include age, gender, and pathological stage, and recalculated the HRs for the 20-, 10-, and 5-gene subsets. All analyses were conducted using the survival package in R.

## Results

### Corelation Analysis

We identified 1,837 genes that were significantly associated with overall survival (OS) time in our univariate Cox regression analysis. To account for multiple testing, false discovery rate (FDR) correction was applied, after which 290 genes remained significant (FDR-adjusted p-value < 0.05). As shown in **Table 1**, the top genes with the strongest positive and negative correlations are presented along with their correlation coefficients (r), raw p-values, and FDR-adjusted p-values. Notably, MAFF (r = 0.25, p = 1.34×10⁻⁹, FDR = 2.46×10⁻⁶) and NR4A3 (r = 0.24, p = 1.26×10⁻⁸, FDR = 9.25×10⁻⁶) exhibited positive associations with OS, whereas LOC728264 (r = –0.21, p = 7.39×10⁻⁷, FDR = 6.36×10⁻⁵) and VAMP1 (r = –0.20, p = 1.20×10⁻⁶, FDR = 0.00013) demonstrated negative associations. A negative correlation indicates that higher expression levels are linked to worse survival outcomes, while a positive correlation suggests that higher expression levels are associated with longer survival. The Pearson correlation coefficient (r) was used to quantify these relationships.

**Table 1.**
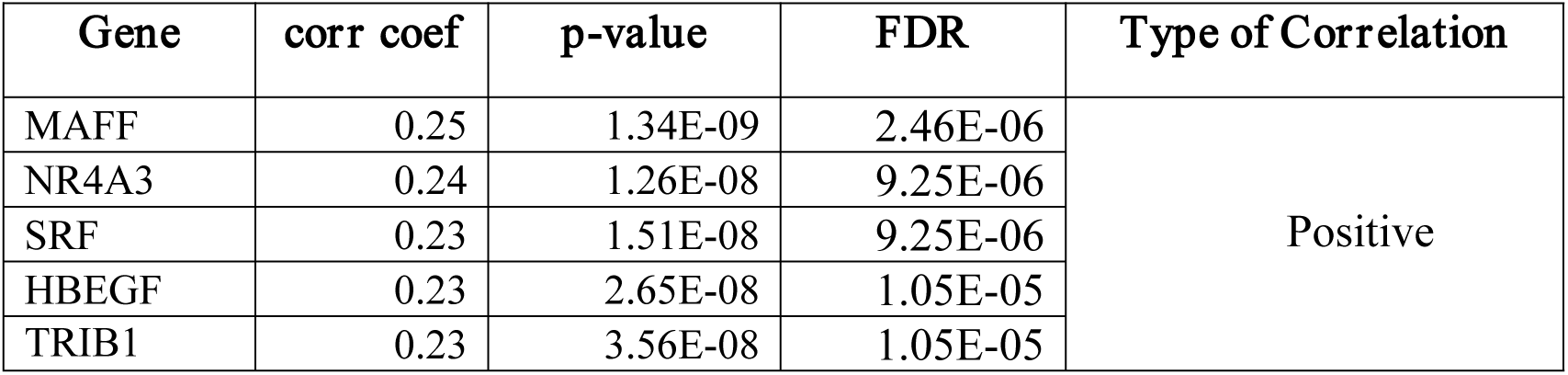
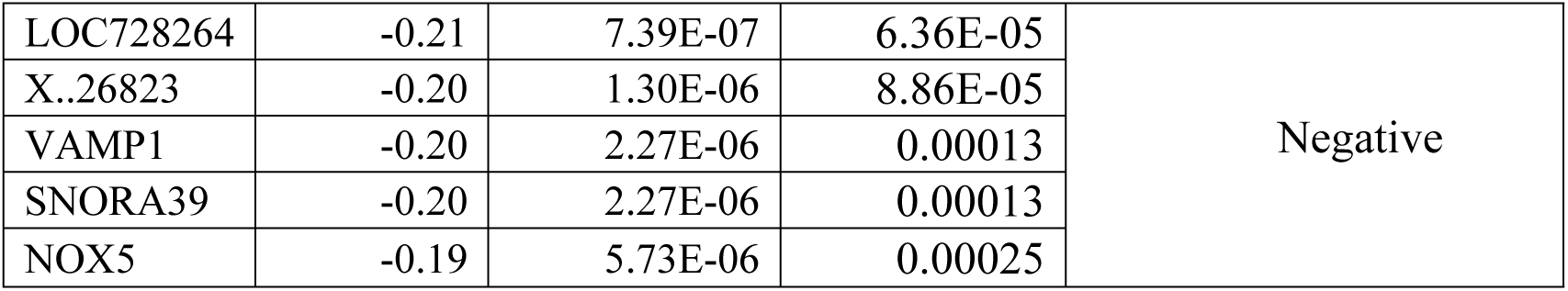
Top Positively and Negatively Correlated Genes are associated with OS Time.

In addition to exploring gene-level associations, we investigated the relationship between key clinical features with OS time. Pearson correlation analysis revealed a weak negative correlation between age and OS time (r = -0.10, p = 0.022), suggesting that increasing age is modestly associated with decreased survival. To further explore this, we compared the mean OS across gender and clinical stages. The mean OS for females was 43.15 months, while males had a lower mean OS of 36.21 months, indicating better survival outcomes in females. Across clinical stages, patients diagnosed at stage II exhibited the highest mean OS of 52.06 months, followed by stage I (41.71 months), stage III (40.14 months), and stage IV (29.59 months), reflecting a clear decline in survival with advancing stage. These findings support the relevance of clinical features, such as age, gender, and stage, in influencing patient prognosis.

We selected 50 positively correlated genes for enrichment analysis. GO analysis identified 965 enriched BP, 51 CC, and 104 MF. In bubble plot, the size of each bubble represents the gene ratio the proportion of genes involved in a given GO term while the color gradient denotes the adjusted p-value, with red indicating stronger statistical significance and blue indicating relatively higher (less significant) p-values. Key enriched terms include “positive regulation of transcription by RNA polymerase II” and “regulation of cell population proliferation”, both showing high gene ratios but appearing in blue, suggesting moderate statistical significance. In contrast, “regulation of cell cycle”, marked in red, exhibits the strongest statistical significance, highlighting its potential importance in thyroid cancer survival as shown in **Figure 3(a)**. Enriched CC terms included “tight junction” and “nucleus,” while MF terms included “sequence-specific DNA binding” and “DNA-binding transcription activator activity, RNA polymerase II-specific” these processes also involved in thyroid cancer. Reactome pathway analysis revealed 221 enriched pathways, Notably, “Signal Transduction (R-HSA-162582)” showed the highest gene ratio and strongest significance, indicating its potential role in thyroid cancer progression shown in **Figure 3(b)**.

**Figure 3.**
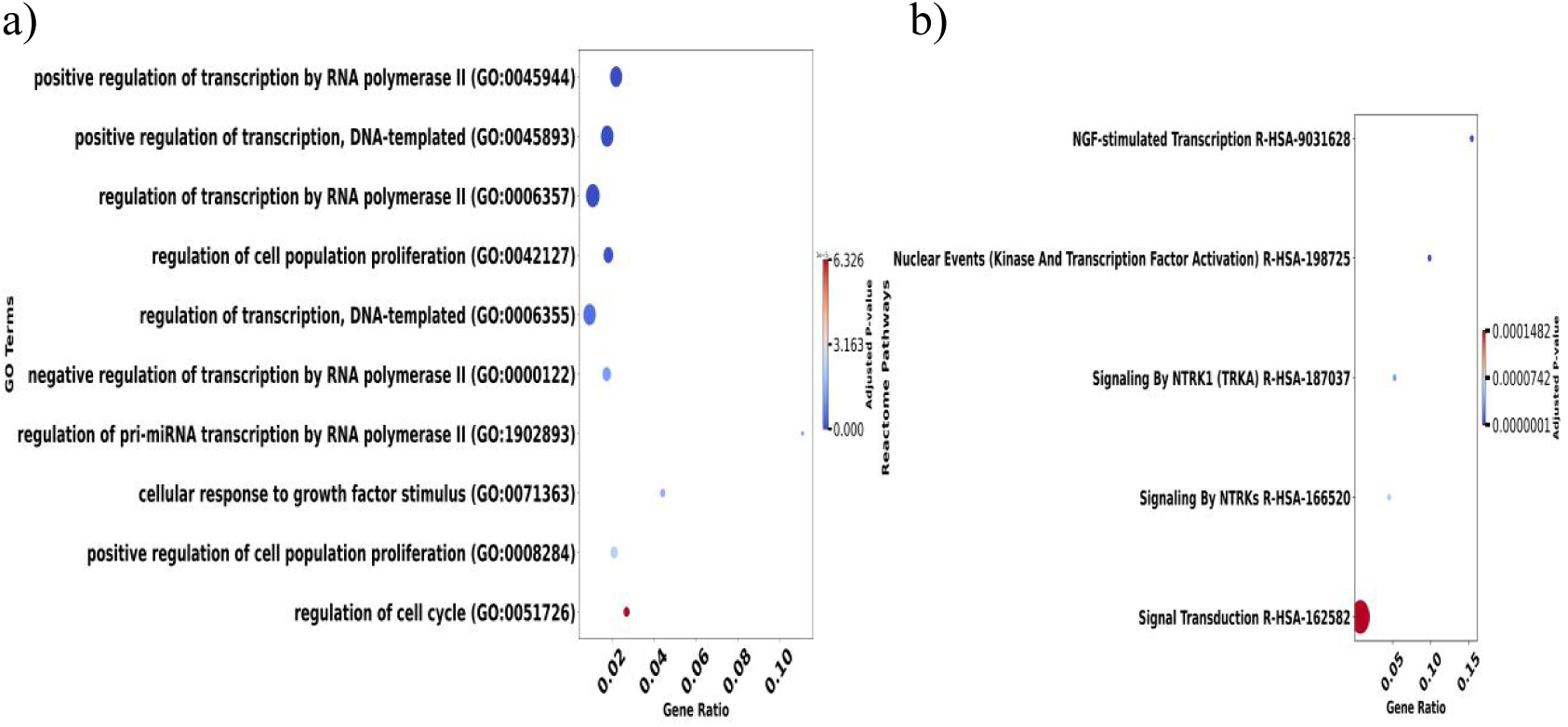
a) Top 10 enriched Gene Ontology biological processes for positively correlated genes b)Top 5 enriched Reactome pathways for positively correlated genes

### Survival Analysis

We performed a regression analysis using univariate Cox proportional hazards. Patients were divided into low-risk (G1) and high-risk (G2) group according to median OS for each gene. The results of this study showed that 883 genes were significantly (p < 0.01) linked to survival outcomes. 541 of these genes had hazard ratios (HR) greater than 1, suggesting a potential link to higher risk, while 342 genes had HR less than 1, suggesting a potential protective function. Based on HR values, the most influential genes are depicted in **Figure 4(a)**, and **Table 2** offers a thorough summary.

**Figure 4.**
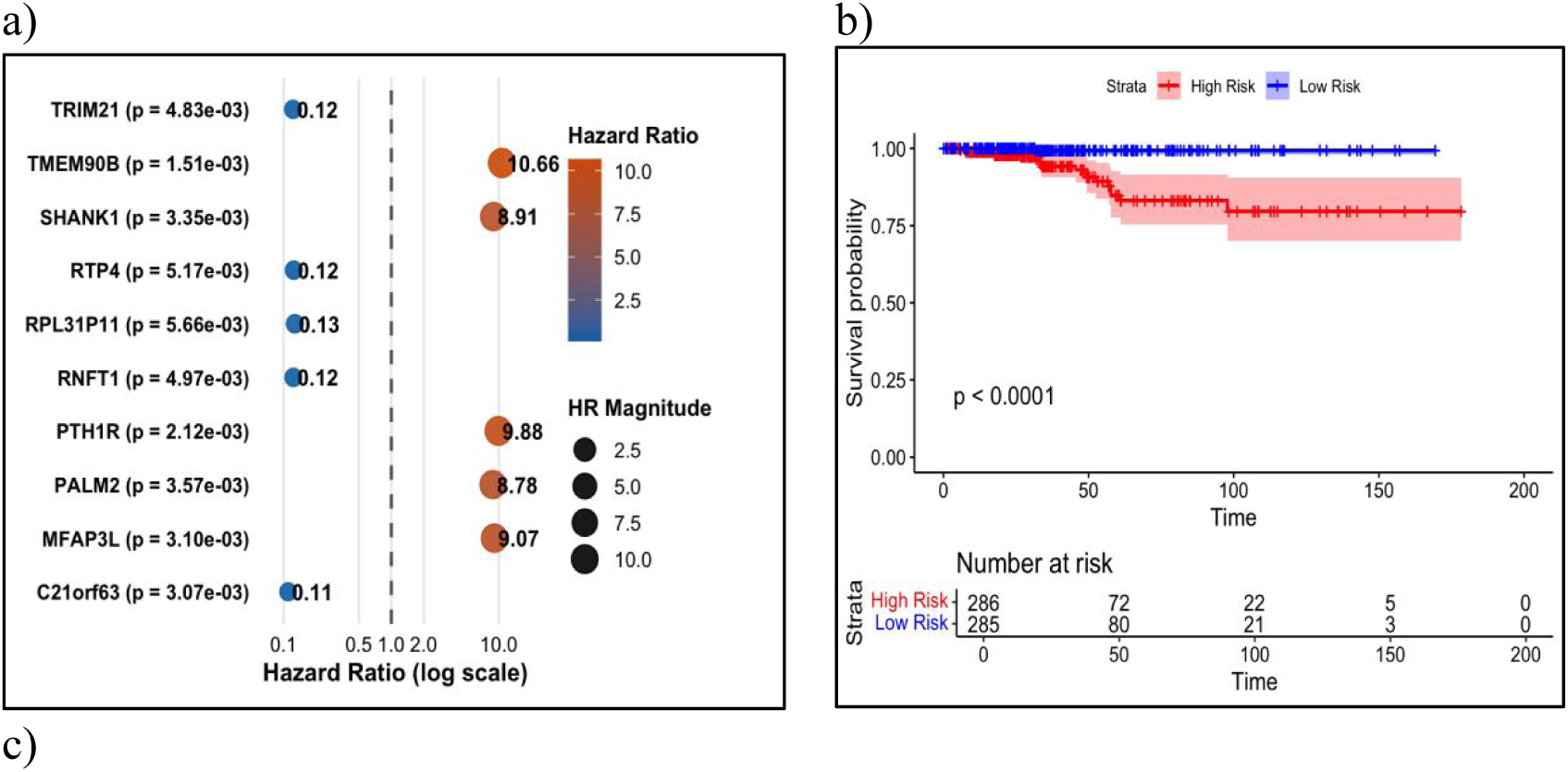
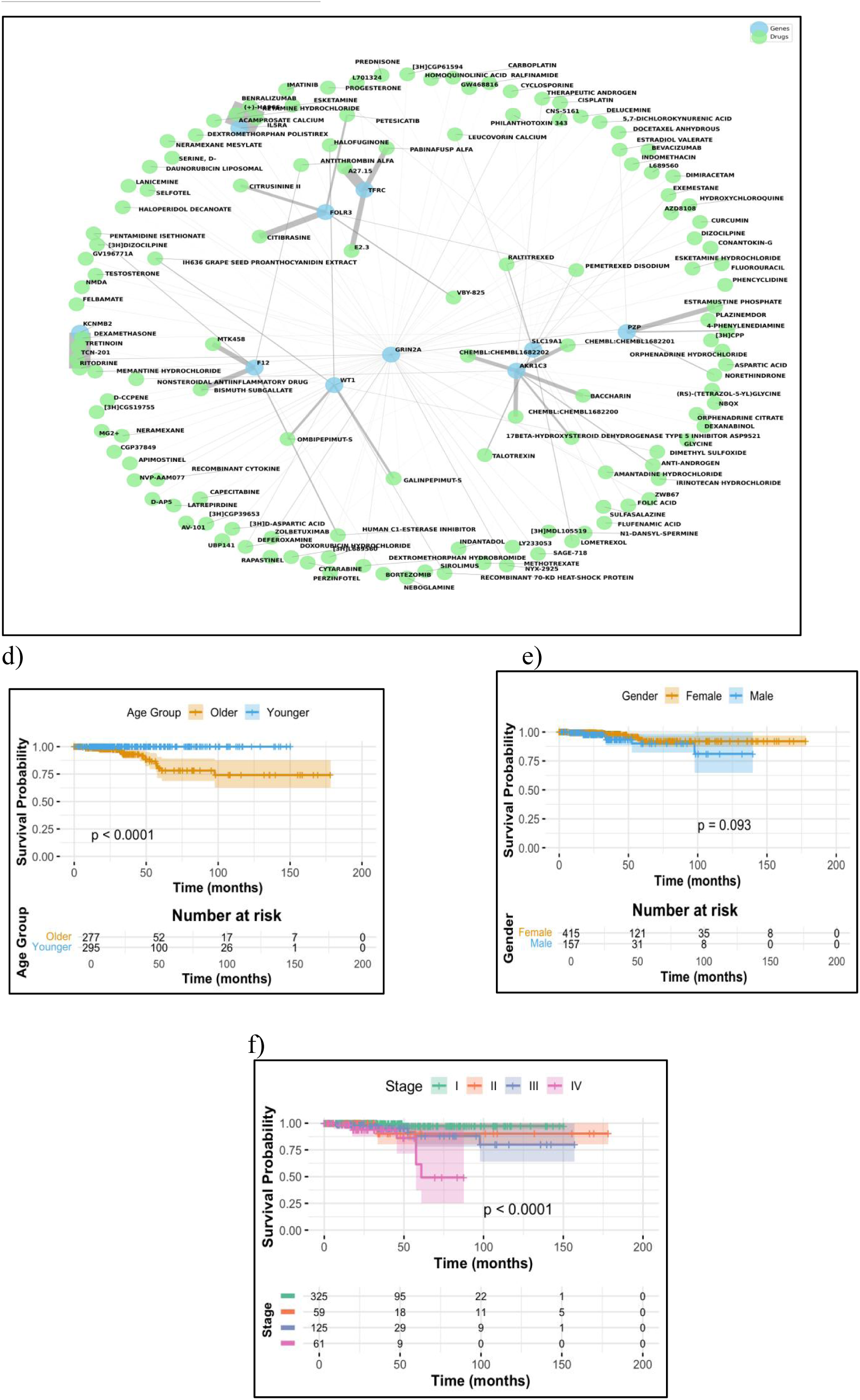
a) Forest plot of hazard ratios with p-values for survival-related genes. b) Kaplan–Meier survival curves comparing overall survival between high- and low-risk groups. c) Gene–Drug Interaction Network illustrating the relationships between 10 genes and their targeting drugs. d) Kaplan–Meier Survival Curve Stratified by Age. e) Kaplan–Meier Survival Curve Stratified by Gender. f) Kaplan–Meier Survival Curve Stratified by Clinical Stage.

**Table 2.**
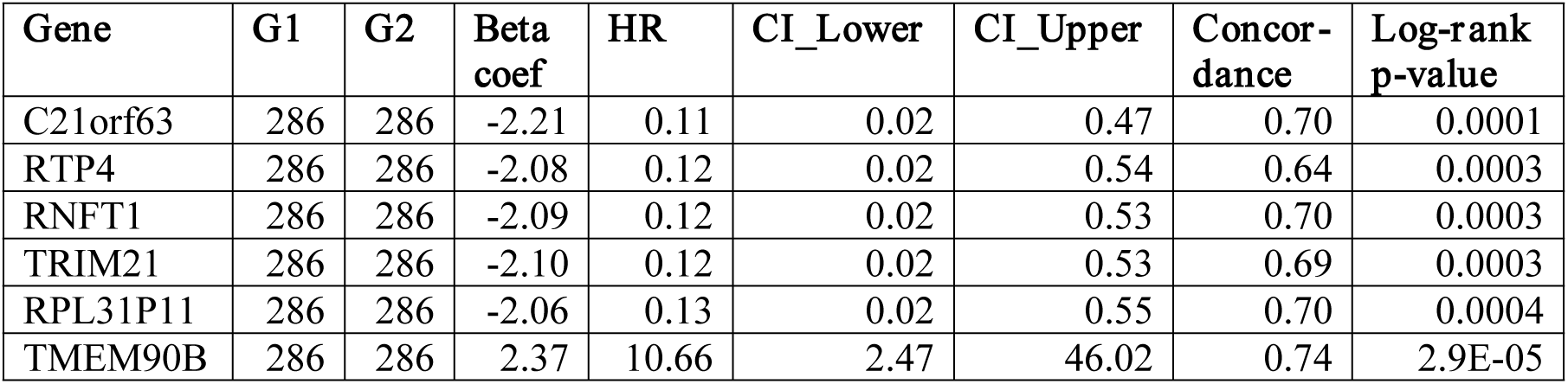
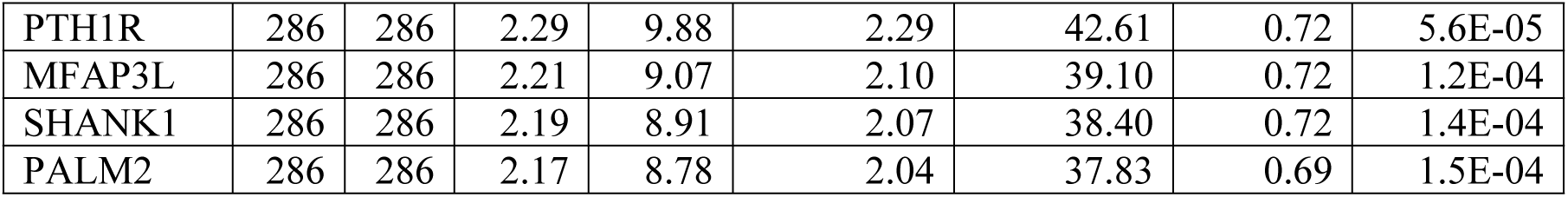
Survival-associated genes with hazard ratios and statistical metrics.

To assess the reproducibility and robustness of previously reported biomarkers, we additionally evaluated nine apoptosis-related genes identified in earlier studies. Notably, all nine genes demonstrated hazard ratios in the same direction as previously reported, confirming consistency across datasets and analytical pipelines. Specifically, ANXA1, CLU, PSEN1, TNFRSF12A, and GPX4 exhibited HR values less than 1, indicating protective effects, whereas TGFBR3, TIMP3, LEF1, and BNIP3L showed HR values greater than 1, suggesting increased risk.

Additionally, age, gender, and clinical stage were analyzed for their impact on overall survival. Age showed a hazard ratio (HR) of 1.11, indicating a slight increase in risk with advancing age. Gender was also significantly associated with survival, with males exhibiting a higher risk of death (HR = 2.12) compared to females. Clinical stage demonstrated a strong association with survival outcomes (HR = 2.50), confirming that higher stages correspond to poorer prognosis. Kaplan–Meier survival curves were plotted for age (using median split) shown in **Figure 4(d)**, gender **Figure 4(e)**, and stage groups **Figure 4(f)**, all of which showed significant separation between groups, further supporting the prognostic relevance of these variables.

We used LASSO Cox regression analysis to identify the most relevant genes linked to overall survival in order to further improve the prognostic model. The optimal penalty parameter (lambda), which was found to be 0.03 based on the smallest cross-validation error, was determined using cross-validation. 31 genes with non-zero coefficients were chosen as a result of this process, demonstrating their high predictive value. As shown in **Table 3**, 13 of these genes served as protective factors with negative beta coef value whereas 17 were linked to increased risk with positive coef value. After that risk score was calculated for each patient using the expression levels and corresponding coefficients of the 31 selected genes. Patients were then stratified into high-risk and low-risk groups based on the median risk score. To evaluate the prognostic value of this stratification, Kaplan-Meier survival analysis was performed shown in **Figure 4(b)**. The results demonstrated a significant difference in overall survival between the two groups, with the high-risk group showing a notably poorer prognosis. Moreover, we found 233 drugs that target 13 of these genes are included in **Supplementary Table S10**. The genes that had the highest drug interaction scores were chosen and they are shown in **Figure 4(c)**.

**Table 3.** List of 31 genes linked to prognosis in THCA.

| Gene | Coef |
| --- | --- |
| STC1 | 0.194 |
| FOLR3 | 0.044 |
| TFRC | 0.022 |
| MYL3 | 0.228 |
| AKR1C3 | 0.068 |
| KCNMB2 | 0.203 |
| SLC19A1 | 0.014 |
| F12 | 0.203 |
| GRIN2A | 0.104 |
| RFPL2 | 0.127 |
| WFIKKN1 | 0.015 |
| DUSP5P | 0.153 |
| TUBAL3 | 0.115 |
| KIAA1024 | 0.016 |
| WT1 | 0.047 |
| PCDHA5 | 0.060 |
| KIAA1644 | 0.164 |
| TUBB2C | -0.330 |
| FLJ35776 | -0.028 |
| X..10357 | -0.426 |
| HTRA4 | -0.058 |
| ASCL2 | -0.100 |
| ZNF367 | -0.023 |
| MRPL42P5 | -0.123 |
| PZP | -0.102 |
| IL5RA | -0.117 |
| HBG1 | -0.069 |
| FLJ44635 | -0.173 |
| REP15 | -0.125 |
| RNF5P1 | -0.018 |
| RPS27 | -0.120 |

### Machine learning Analysis

To identify the most relevant genomic and epigenomic markers, we used a variety of well-established feature-selection methods, such as SVC-L1, SelectKBest, RFE refer to **Supplementary Tables (S1–S9**). Several machine-learning classifiers, including DT, LightGBM, SVM, ExtraTrees (39, 40), were trained using these candidate biomarkers. The accuracy, AUC, specificity, sensitivity, and Matthews correlation coefficient (MCC) of the models were utilized to assess their performance. On the primary set of 20 prognostic genes we achieved an AUC 0.93 with MCC value 0.74 shown in **Table 5**.

**Table 5.**
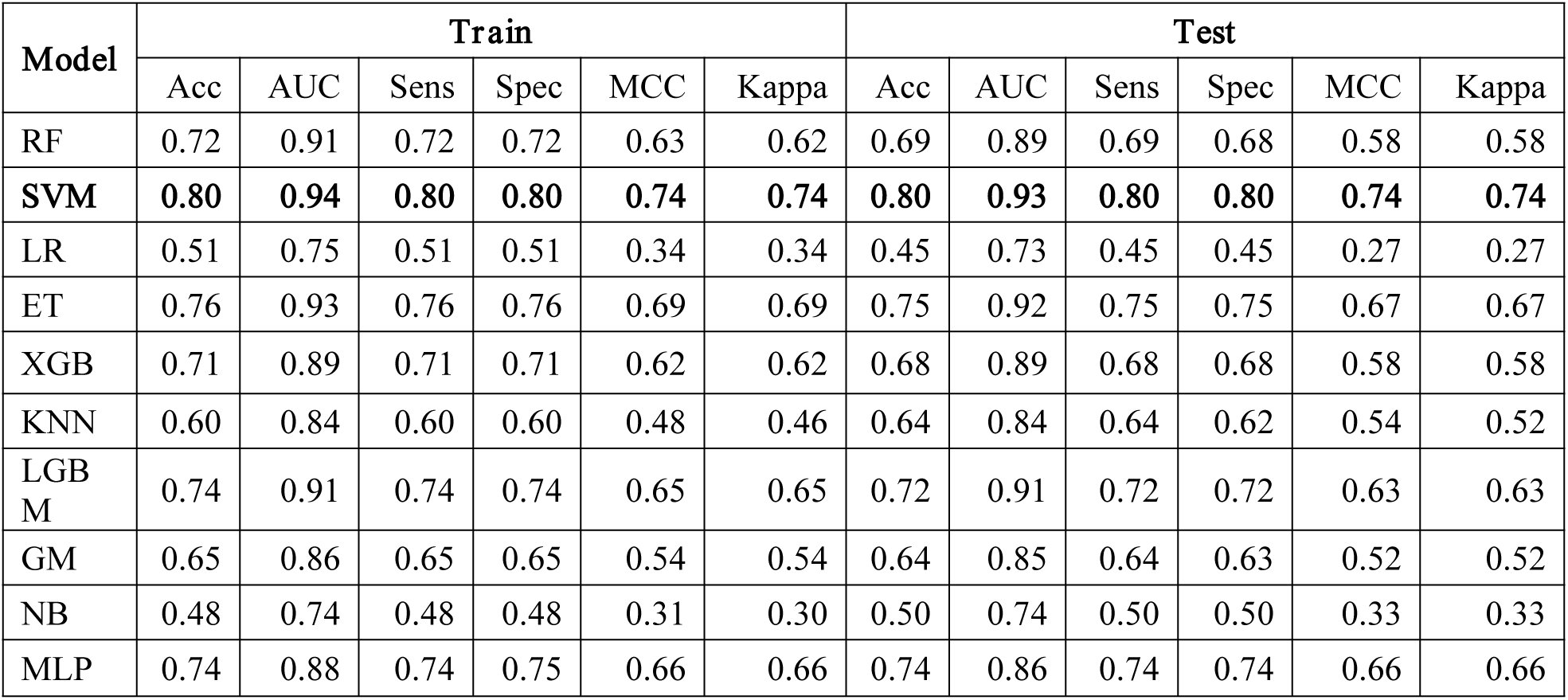
Performance measures of the 20 primary genes selected through SVC-L1 for classification of survival categories in train and test datasets.

Similarly, SVM achieved the highest performance as compare to all the models, with an MCC of 0.76 and an AUC of 0.94 on the second set of prognostic biomarkers. Additionally on the third set we achieved AUC of 0.91. The fourth (AUC: 0.93), fifth (AUC:0.93), sixth (AUC: 0.92), and seventh (AUC: 0.92) biomarker sets all showed consistent performance shown in **Table 6**. Moreover, we achieved an AUC of 0.94 and a Kappa of 0.70 on the test dataset by employing the SVC-L1 technique to select 20 genes from a total of seven biomarker sets.

**Table 6.** Performance measures for a distinct set of biomarkers selected through SVC-L1.

| For the second set of biomarkers |  |  |  |  |  |  |  |  |  |  |  |  |
| --- | --- | --- | --- | --- | --- | --- | --- | --- | --- | --- | --- | --- |
| Model | Train |  |  |  |  |  | Test |  |  |  |  |  |
|  | Acc | AUC | Sens | Spec | MCC | Kappa | Acc | AUC | Sens | Spec | MCC | Kappa |
| RF | 0.74 | 0.91 | 0.74 | 0.74 | 0.66 | 0.66 | 0.75 | 0.92 | 0.75 | 0.74 | 0.67 | 0.66 |
| **SVM** | **0.79** | **0.94** | **0.79** | **0.79** | **0.72** | **0.72** | **0.82** | **0.94** | **0.82** | **0.81** | **0.76** | **0.76** |
| LR | 0.46 | 0.73 | 0.46 | 0.46 | 0.28 | 0.28 | 0.41 | 0.70 | 0.41 | 0.41 | 0.21 | 0.21 |
| ET | 0.79 | 0.94 | 0.79 | 0.79 | 0.73 | 0.72 | 0.81 | 0.94 | 0.81 | 0.81 | 0.75 | 0.75 |
| XGB | 0.71 | 0.90 | 0.71 | 0.71 | 0.62 | 0.62 | 0.74 | 0.91 | 0.74 | 0.73 | 0.65 | 0.65 |
| KNN | 0.61 | 0.83 | 0.61 | 0.62 | 0.49 | 0.48 | 0.60 | 0.84 | 0.60 | 0.58 | 0.48 | 0.47 |
| LGBM | 0.75 | 0.92 | 0.75 | 0.75 | 0.67 | 0.67 | 0.77 | 0.93 | 0.77 | 0.77 | 0.69 | 0.69 |
| GM | 0.68 | 0.88 | 0.68 | 0.68 | 0.57 | 0.57 | 0.67 | 0.90 | 0.67 | 0.66 | 0.56 | 0.55 |
| NB | 0.43 | 0.70 | 0.43 | 0.43 | 0.24 | 0.24 | 0.39 | 0.67 | 0.39 | 0.38 | 0.20 | 0.19 |
| MLP | 0.73 | 0.87 | 0.73 | 0.72 | 0.64 | 0.63 | 0.77 | 0.88 | 0.77 | 0.76 | 0.69 | 0.69 |
| For the third set of biomarkers |  |  |  |  |  |  |  |  |  |  |  |  |
| RF | 0.70 | 0.90 | 0.70 | 0.70 | 0.61 | 0.61 | 0.69 | 0.89 | 0.69 | 0.68 | 0.59 | 0.58 |
| **SVM** | **0.77** | **0.94** | **0.77** | **0.77** | **0.70** | **0.70** | **0.76** | **0.91** | **0.76** | **0.76** | **0.69** | **0.68** |
| LR | 0.49 | 0.74 | 0.49 | 0.49 | 0.32 | 0.31 | 0.38 | 0.66 | 0.38 | 0.37 | 0.17 | 0.17 |
| **ET** | **0.76** | **0.93** | **0.76** | **0.76** | **0.69** | **0.68** | **0.77** | **0.92** | **0.77** | **0.75** | **0.69** | **0.69** |
| XGB | 0.69 | 0.88 | 0.69 | 0.69 | 0.58 | 0.58 | 0.71 | 0.88 | 0.71 | 0.70 | 0.62 | 0.61 |
| KNN | 0.57 | 0.82 | 0.57 | 0.60 | 0.44 | 0.43 | 0.59 | 0.82 | 0.59 | 0.57 | 0.46 | 0.45 |
| LGBM | 0.73 | 0.91 | 0.73 | 0.73 | 0.64 | 0.64 | 0.73 | 0.90 | 0.73 | 0.72 | 0.65 | 0.65 |
| GM | 0.68 | 0.88 | 0.68 | 0.68 | 0.57 | 0.57 | 0.75 | 0.92 | 0.75 | 0.75 | 0.67 | 0.67 |
| NB | 0.46 | 0.72 | 0.46 | 0.46 | 0.29 | 0.28 | 0.35 | 0.64 | 0.35 | 0.33 | 0.14 | 0.14 |
| MLP | 0.73 | 0.87 | 0.73 | 0.72 | 0.64 | 0.63 | 0.70 | 0.83 | 0.70 | 0.69 | 0.60 | 0.60 |
| For the Fourth set of biomarkers |  |  |  |  |  |  |  |  |  |  |  |  |
| RF | 0.71 | 0.90 | 0.71 | 0.71 | 0.62 | 0.62 | 0.76 | 0.90 | 0.76 | 0.76 | 0.69 | 0.68 |
| **SVM** | **0.80** | **0.94** | **0.80** | **0.80** | **0.74** | **0.73** | **0.79** | **0.93** | **0.79** | **0.78** | **0.72** | **0.72** |
| LR | 0.51 | 0.76 | 0.51 | 0.51 | 0.35 | 0.35 | 0.43 | 0.70 | 0.43 | 0.42 | 0.24 | 0.24 |
| ET | 0.77 | 0.93 | 0.77 | 0.77 | 0.69 | 0.69 | 0.77 | 0.93 | 0.77 | 0.76 | 0.70 | 0.69 |
| XGB | 0.70 | 0.90 | 0.70 | 0.70 | 0.60 | 0.60 | 0.73 | 0.90 | 0.73 | 0.72 | 0.65 | 0.65 |
| KNN | 0.64 | 0.85 | 0.64 | 0.63 | 0.53 | 0.51 | 0.65 | 0.85 | 0.65 | 0.59 | 0.55 | 0.53 |
| LGBM | 0.74 | 0.92 | 0.74 | 0.73 | 0.65 | 0.65 | 0.78 | 0.92 | 0.78 | 0.78 | 0.71 | 0.71 |
| GM | 0.66 | 0.86 | 0.66 | 0.65 | 0.54 | 0.54 | 0.71 | 0.91 | 0.71 | 0.70 | 0.61 | 0.61 |
| NB | 0.48 | 0.73 | 0.48 | 0.48 | 0.31 | 0.30 | 0.46 | 0.69 | 0.46 | 0.46 | 0.28 | 0.28 |
| MLP | 0.76 | 0.89 | 0.76 | 0.76 | 0.68 | 0.68 | 0.77 | 0.86 | 0.77 | 0.76 | 0.69 | 0.69 |
| For the fifth set of biomarkers |  |  |  |  |  |  |  |  |  |  |  |  |
| RF | 0.73 | 0.90 | 0.73 | 0.73 | 0.63 | 0.63 | 0.69 | 0.90 | 0.69 | 0.69 | 0.59 | 0.59 |
| **SVM** | **0.80** | **0.93** | **0.80** | **0.79** | **0.73** | **0.73** | **0.76** | **0.93** | **0.76** | **0.74** | **0.68** | **0.68** |
| LR | 0.46 | 0.72 | 0.46 | 0.46 | 0.28 | 0.28 | 0.41 | 0.67 | 0.41 | 0.42 | 0.22 | 0.22 |
| **ET** | **0.78** | **0.93** | **0.78** | **0.78** | **0.71** | **0.70** | **0.76** | **0.94** | **0.76** | **0.75** | **0.69** | **0.68** |
| XGB | 0.71 | 0.89 | 0.71 | 0.70 | 0.61 | 0.61 | 0.68 | 0.88 | 0.68 | 0.68 | 0.57 | 0.57 |
| KNN | 0.59 | 0.83 | 0.59 | 0.60 | 0.47 | 0.45 | 0.63 | 0.85 | 0.63 | 0.62 | 0.52 | 0.50 |
| LGBM | 0.73 | 0.92 | 0.73 | 0.73 | 0.65 | 0.65 | 0.72 | 0.91 | 0.72 | 0.72 | 0.63 | 0.63 |
| GM | 0.72 | 0.92 | 0.72 | 0.71 | 0.62 | 0.62 | 0.74 | 0.92 | 0.74 | 0.74 | 0.65 | 0.65 |
| NB | 0.40 | 0.68 | 0.40 | 0.43 | 0.21 | 0.20 | 0.39 | 0.65 | 0.39 | 0.40 | 0.19 | 0.18 |
| MLP | 0.74 | 0.89 | 0.74 | 0.74 | 0.66 | 0.66 | 0.72 | 0.87 | 0.72 | 0.71 | 0.63 | 0.63 |

|  | For the Sixth set of biomarkers |  |  |  |  |  |  |  |  |  |  |  |
| --- | --- | --- | --- | --- | --- | --- | --- | --- | --- | --- | --- | --- |
| RF | 0.70 | 0.89 | 0.70 | 0.70 | 0.61 | 0.60 | 0.74 | 0.91 | 0.74 | 0.73 | 0.65 | 0.65 |
| **SVM** | **0.75** | **0.92** | **0.75** | **0.75** | **0.67** | **0.67** | **0.77** | **0.92** | **0.77** | **0.76** | **0.69** | **0.69** |
| LR | 0.47 | 0.70 | 0.47 | 0.48 | 0.30 | 0.30 | 0.49 | 0.69 | 0.49 | 0.49 | 0.32 | 0.32 |
| **ET** | **0.76** | **0.92** | **0.76** | **0.76** | **0.68** | **0.68** | **0.77** | **0.94** | **0.77** | **0.77** | **0.70** | **0.69** |
| XGB | 0.67 | 0.88 | 0.67 | 0.67 | 0.56 | 0.56 | 0.68 | 0.89 | 0.68 | 0.68 | 0.58 | 0.58 |
| KNN | 0.56 | 0.80 | 0.56 | 0.57 | 0.43 | 0.41 | 0.66 | 0.87 | 0.66 | 0.68 | 0.57 | 0.55 |
| LGBM | 0.72 | 0.90 | 0.72 | 0.72 | 0.62 | 0.62 | 0.73 | 0.91 | 0.73 | 0.73 | 0.65 | 0.65 |
| GM | 0.68 | 0.88 | 0.68 | 0.68 | 0.57 | 0.57 | 0.71 | 0.90 | 0.71 | 0.70 | 0.61 | 0.61 |
| NB | 0.43 | 0.68 | 0.43 | 0.43 | 0.24 | 0.24 | 0.44 | 0.70 | 0.44 | 0.44 | 0.26 | 0.26 |
| MLP | 0.70 | 0.84 | 0.70 | 0.70 | 0.61 | 0.60 | 0.67 | 0.84 | 0.67 | 0.65 | 0.56 | 0.55 |
|  | For the Seventh set of biomarkers |  |  |  |  |  |  |  |  |  |  |  |
| RF | 0.69 | 0.90 | 0.69 | 0.69 | 0.59 | 0.59 | 0.70 | 0.89 | 0.70 | 0.71 | 0.61 | 0.60 |
| **SVM** | **0.78** | **0.93** | **0.78** | **0.78** | **0.71** | **0.71** | **0.76** | **0.92** | **0.76** | **0.76** | **0.68** | **0.68** |
| LR | 0.50 | 0.73 | 0.50 | 0.49 | 0.33 | 0.33 | 0.45 | 0.71 | 0.45 | 0.45 | 0.27 | 0.27 |
| ET | 0.75 | 0.93 | 0.75 | 0.75 | 0.67 | 0.67 | 0.73 | 0.92 | 0.73 | 0.73 | 0.65 | 0.65 |
| XGB | 0.69 | 0.88 | 0.69 | 0.69 | 0.59 | 0.59 | 0.67 | 0.88 | 0.67 | 0.68 | 0.56 | 0.56 |
| KNN | 0.57 | 0.81 | 0.57 | 0.57 | 0.44 | 0.43 | 0.60 | 0.85 | 0.60 | 0.65 | 0.49 | 0.47 |
| LGBM | 0.71 | 0.91 | 0.71 | 0.71 | 0.62 | 0.62 | 0.71 | 0.90 | 0.71 | 0.71 | 0.62 | 0.61 |
| GM | 0.63 | 0.86 | 0.63 | 0.62 | 0.51 | 0.51 | 0.72 | 0.90 | 0.72 | 0.72 | 0.62 | 0.62 |
| NB | 0.49 | 0.72 | 0.49 | 0.48 | 0.32 | 0.31 | 0.51 | 0.73 | 0.51 | 0.51 | 0.35 | 0.35 |
| MLP | 0.72 | 0.87 | 0.72 | 0.71 | 0.63 | 0.63 | 0.68 | 0.85 | 0.68 | 0.69 | 0.58 | 0.58 |

To assess our best-performing model’s (SVM) performance using AUC scores for four classes over seven different biomarker sets: Primary, Secondary, Third, Fourth, Fifth, Sixth, and Seventh. All biomarker sets demonstrated good predictive powers, according to the research, with AUC values continuously over 0.85 for the majority of classes, highlighting the dependability and robustness of the chosen features. The Secondary biomarker set, which obtained AUCs of 0.99, 0.90, 0.91, and 0.95 for Classes 0 through 3, respectively, showed the most consistent and balanced performance across all four classes shwon in **Table 7**. This indicates its strong potential as a generalized predictor across patient subtypes. Nonetheless, certain biomarker sets also demonstrated better results for distinct classes. With the Third and Fifth sets, for instance, Class 0 achieved a perfect AUC of 1.00; the Primary and Secondary sets predicted the best for Class 1 (AUC = 0.90); the Fourth set did the best for Class 2 (AUC = 0.92); and the Fourth set produced the highest AUC of 0.98 for Class 3. These results suggest that while the Secondary set offers strong generalizability, other biomarker sets may capture unique class-specific biological signatures.

**Table 7.** AUC for each class using our Best performing Model(SVM) in all seven sets of Biomarkers.

| Class | Primary | Secondary | Third | Fourth | Fifth | Sixth | Seventh |
| --- | --- | --- | --- | --- | --- | --- | --- |
| 0 | 0.97 | 0.99 | 1.00 | 0.98 | 1.00 | 0.99 | 0.96 |
| 1 | 0.90 | 0.90 | 0.83 | 0.85 | 0.85 | 0.86 | 0.88 |
| 2 | 0.91 | 0.91 | 0.90 | 0.92 | 0.91 | 0.89 | 0.89 |
| 3 | 0.93 | 0.95 | 0.93 | 0.98 | 0.96 | 0.94 | 0.96 |

### Ensemble Models

To enhance the predictive performance, ensemble learning methods were applied using the 20 prognostic genes identified from the primary biomarker set. A Voting classifier, integrating RF, ET, LightGBM, and SVM, produced an AUC of 0.93 and a Matthews Correlation Coefficient (MCC) of 0.70, reflecting robust classification ability. Additionally, Stacking classifiers were explored with two configurations: one using SVM as the meta-learner with RF, ET, and LightGBM as base models. This stacking approach yielded strong results, achieving an AUC of 0.93 and an MCC of 0.68.

For the secondary biomarker set, ensemble methods demonstrated comparable and even slightly improved performance. The Voting classifier using the same base models achieved an AUC of 0.95 and an MCC of 0.72. Similarly, the Stacking classifier with SVM as the meta-learner attained an AUC of 0.95 and an MCC of 0.71, highlighting the effectiveness of the secondary gene set in capturing prognostic signals. These findings emphasize the strength of ensemble strategies and the value of different biomarker sets in improving classification performance. A detailed comparison of these ensemble strategies is presented in **Table 8**.

**Table 8.** Performance on Primary and secondary set of biomarkers using Ensemble model.

| Model | Combination of RF,ET,LGBM & SVM for primary set |  |  |  |  |  |  |  |  |  |  |  |
| --- | --- | --- | --- | --- | --- | --- | --- | --- | --- | --- | --- | --- |
|  | Train |  |  |  |  |  | Test |  |  |  |  |  |
|  | Acc | AUC | Sens | Spec | MCC | Kappa | Acc | AUC | Sens | Spec | MCC | Kappa |
| Voting | 0.78 | 0.94 | 0.78 | 0.78 | 0.70 | 0.70 | 0.78 | 0.93 | 0.78 | 0.77 | 0.70 | 0.70 |
| SVM as Meta-model & RF,ET,LGBM as base classifiers |  |  |  |  |  |  |  |  |  |  |  |  |
| Stacking | 0.76 | 0.93 | 0.76 | 0.76 | 0.67 | 0.67 | 0.76 | 0.93 | 0.76 | 0.76 | 0.68 | 0.68 |
| RF,ET,LGBM & SVM for Secondary set |  |  |  |  |  |  |  |  |  |  |  |  |
| Voting | 0.79 | 0.94 | 0.79 | 0.79 | 0.72 | 0.71 | 0.79 | 0.95 | 0.79 | 0.79 | 0.73 | 0.72 |
| SVM as meta model & RF,ET,LGBM as base classifiers |  |  |  |  |  |  |  |  |  |  |  |  |
| Stacking | 0.76 | 0.94 | 0.76 | 0.76 | 0.68 | 0.68 | 0.78 | 0.95 | 0.78 | 0.78 | 0.71 | 0.71 |

Overall, our analysis showed consistently high classification accuracy across a variety of gene sets, with all seven biomarker sets achieving AUC values above 0.91. These outcomes highlight how effective and dependable our method is at finding highly predictive prognostic biomarkers.

### Prognostic Efficiency of reduced subset Across Biomarker Sets

The predictive performance of the top 10 genes from each of the seven biomarker sets was evaluated using both training and test datasets shown in **Table 9** with the ET classifier. Across the training sets, AUC values ranged from 0.86 to 0.90, with MCC values between 0.55 and 0.64, indicating strong discriminative ability and balanced classification. On the test sets, AUCs remained high (0.87–0.90), while MCC values ranged from 0.54 to 0.61, confirming reliable performance on unseen data. Among the seven sets, the 5th biomarker set demonstrated the best overall performance, achieving the highest training AUC (0.90) and MCC (0.64), with strong test AUC (0.90). The 4th set showed slightly higher test MCC (0.61) but lower training MCC (0.59). Overall, the top 10-genes, especially the 5th set, performed well. They showed strong prognostic ability and were almost as effective as the full 20-gene biomarker sets.

**Table 9.** Performance measures of the top 10 genes on all the seven sets.

| Set | Train |  |  |  |  |  | Test |  |  |  |  |  |
| --- | --- | --- | --- | --- | --- | --- | --- | --- | --- | --- | --- | --- |
|  | Acc | AUC | Sens | Spec | MCC | Kappa | Acc | AUC | Sens | Spec | MCC | Kappa |
| 1st | 0.69 | 0.89 | 0.69 | 0.69 | 0.59 | 0.58 | 0.69 | 0.87 | 0.69 | 0.69 | 0.58 | 0.58 |
| 2nd | 0.67 | 0.86 | 0.67 | 0.67 | 0.56 | 0.56 | 0.66 | 0.89 | 0.66 | 0.66 | 0.54 | 0.54 |
| 3rd | 0.69 | 0.89 | 0.69 | 0.69 | 0.59 | 0.59 | 0.67 | 0.89 | 0.67 | 0.67 | 0.56 | 0.56 |
| 4th | 0.69 | 0.89 | 0.69 | 0.69 | 0.59 | 0.59 | 0.71 | 0.89 | 0.71 | 0.69 | 0.61 | 0.61 |
| **5th** | **0.73** | **0.90** | **0.73** | **0.73** | **0.64** | **0.64** | **0.68** | **0.90** | **0.68** | **0.68** | **0.58** | **0.58** |
| 6th | 0.70 | 0.89 | 0.70 | 0.71 | 0.61 | 0.60 | 0.69 | 0.89 | 0.69 | 0.69 | 0.59 | 0.59 |
| 7th | 0.66 | 0.87 | 0.66 | 0.65 | 0.55 | 0.55 | 0.67 | 0.89 | 0.67 | 0.66 | 0.56 | 0.55 |

The top 5 genes from each biomarker set showed moderate but consistent predictive performance shown in **Table 10**. Training AUCs ranged from 0.78 to 0.82 and MCCs from 0.41 to 0.47, while test AUCs were 0.78–0.86 and MCCs 0.40–0.51. The 2nd panel performed best, with the highest test AUC (0.86) and MCC (0.51). Overall, even small 5-gene panels provided good prognostic predictions, though slightly lower than the 10- or 20-gene sets.

**Table 10.** Performance measures of the top 5 genes on all the seven sets.

| Set | Train |  |  |  |  |  | Test |  |  |  |  |  |
| --- | --- | --- | --- | --- | --- | --- | --- | --- | --- | --- | --- | --- |
|  | Acc | AUC | Sens | Spec | MCC | Kappa | Acc | AUC | Sens | Spec | MCC | Kappa |
| 1st | 0.58 | 0.81 | 0.58 | 0.58 | 0.44 | 0.44 | 0.55 | 0.79 | 0.55 | 0.55 | 0.40 | 0.39 |
| **2nd** | **0.60** | **0.82** | **0.60** | **0.59** | **0.47** | **0.47** | **0.63** | **0.86** | **0.63** | **0.63** | **0.51** | **0.50** |
| 3rd | 0.55 | 0.78 | 0.55 | 0.54 | 0.41 | 0.41 | 0.56 | 0.80 | 0.56 | 0.56 | 0.42 | 0.42 |
| 4th | 0.56 | 0.79 | 0.56 | 0.56 | 0.41 | 0.41 | 0.58 | 0.81 | 0.58 | 0.57 | 0.45 | 0.44 |
| 5th | 0.59 | 0.82 | 0.59 | 0.60 | 0.46 | 0.45 | 0.56 | 0.78 | 0.56 | 0.57 | 0.42 | 0.42 |
| 6th | 0.56 | 0.80 | 0.56 | 0.55 | 0.41 | 0.41 | 0.56 | 0.79 | 0.56 | 0.56 | 0.42 | 0.42 |
| 7th | 0.57 | 0.80 | 0.57 | 0.57 | 0.44 | 0.43 | 0.58 | 0.82 | 0.58 | 0.57 | 0.44 | 0.44 |

### Integration of Clinical Features

Integrating clinical variables (age, gender, and stage) with the primary biomarker set significantly improved model performance. The best-performing model, SVM achieved an AUC of 0.94 on the training set and 0.96 on the test set, along with a Kappa score of 0.80 shown in **Table 11**. For comparison, a model trained using only the clinical features (age, gender, and stage) achieved an AUC of 0.76 on the test set using the LGBM highlighting the added predictive value of the gene expression biomarkers when combined with clinical data.

**Table 11.** Performance of Machine Learning Models on the primary Biomarker Set with Clinical Features.

| Model | Train |  |  |  |  |  | Test |  |  |  |  |  |
| --- | --- | --- | --- | --- | --- | --- | --- | --- | --- | --- | --- | --- |
|  | Acc | AUC | Sens | Spec | MCC | Kappa | Acc | AUC | Sens | Spec | MCC | Kappa |
| RF | 0.75 | 0.92 | 0.75 | 0.75 | 0.67 | 0.67 | 0.78 | 0.93 | 0.78 | 0.77 | 0.70 | 0.70 |
| **SVM** | **0.79** | **0.94** | **0.79** | **0.79** | **0.72** | **0.72** | **0.85** | **0.96** | **0.85** | **0.85** | **0.80** | **0.80** |
| ET | 0.79 | 0.94 | 0.79 | 0.79 | 0.72 | 0.72 | 0.83 | 0.95 | 0.83 | 0.83 | 0.77 | 0.77 |
| XGB | 0.72 | 0.91 | 0.72 | 0.72 | 0.63 | 0.63 | 0.80 | 0.92 | 0.80 | 0.80 | 0.73 | 0.73 |
| KNN | 0.60 | 0.84 | 0.60 | 0.62 | 0.48 | 0.46 | 0.63 | 0.83 | 0.63 | 0.65 | 0.52 | 0.50 |
| LGBM | 0.76 | 0.93 | 0.76 | 0.76 | 0.69 | 0.68 | 0.79 | 0.94 | 0.79 | 0.78 | 0.72 | 0.72 |
| GM | 0.70 | 0.90 | 0.70 | 0.70 | 0.60 | 0.60 | 0.72 | 0.92 | 0.72 | 0.71 | 0.63 | 0.63 |
| NB | 0.46 | 0.73 | 0.46 | 0.46 | 0.28 | 0.28 | 0.47 | 0.73 | 0.47 | 0.47 | 0.30 | 0.30 |
| MLP | 0.74 | 0.89 | 0.74 | 0.74 | 0.66 | 0.66 | 0.77 | 0.91 | 0.77 | 0.77 | 0.69 | 0.69 |

For the primary biomarker set, the model demonstrated strong discriminatory power across all survival classes. The AUC values for Class 0 (0–1 year), Class 1 (1–3 years), Class 2 (3–5 years), and Class 3 (>5 years) were 0.99, 0.91, 0.96, and 0.95, respectively shown in **Figure 5**. These high AUC scores highlight the model’s ability to accurately distinguish patients across different survival risk categories.

**Figure 5.**
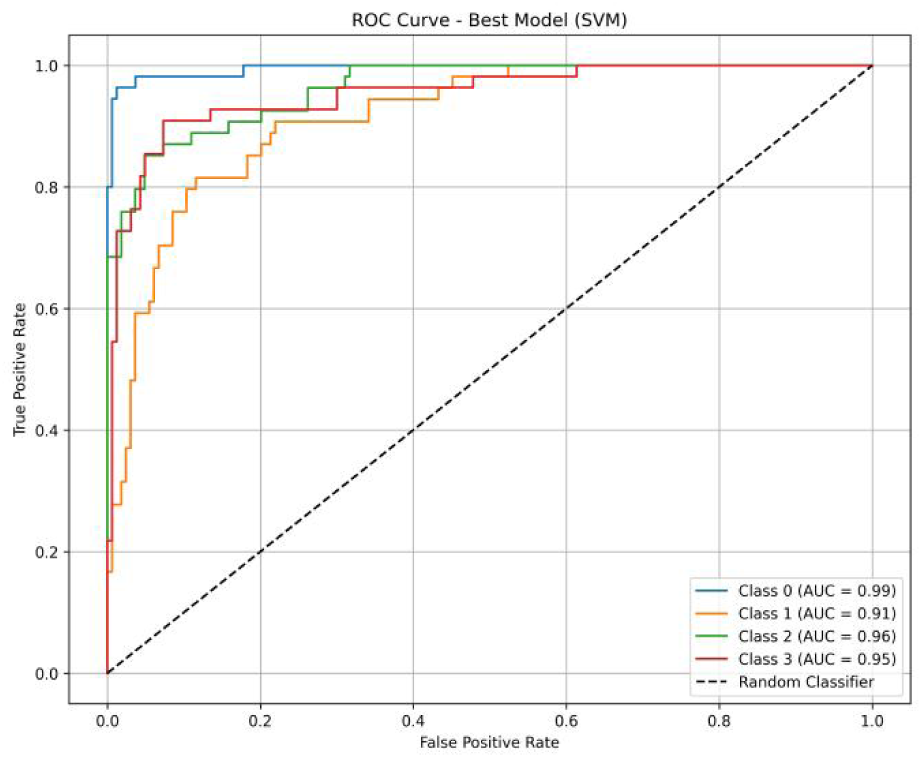
ROC curves showing high AUCs across all survival classes using the primary biomarker set along with clinical features

Second biomarker set with clinical features led to robust model performance. The best model achieved an AUC of 0.94 on the training set and 0.95 on the test set, with a Kappa score of 0.76 depicted in **Table 12** indicating strong predictive ability and consistency in distinguishing patient risk groups. Notably, when clinical factors were added, the performance of the 20-gene set improved, whereas the 10- and 5-gene set remained same.

**Table 12.**
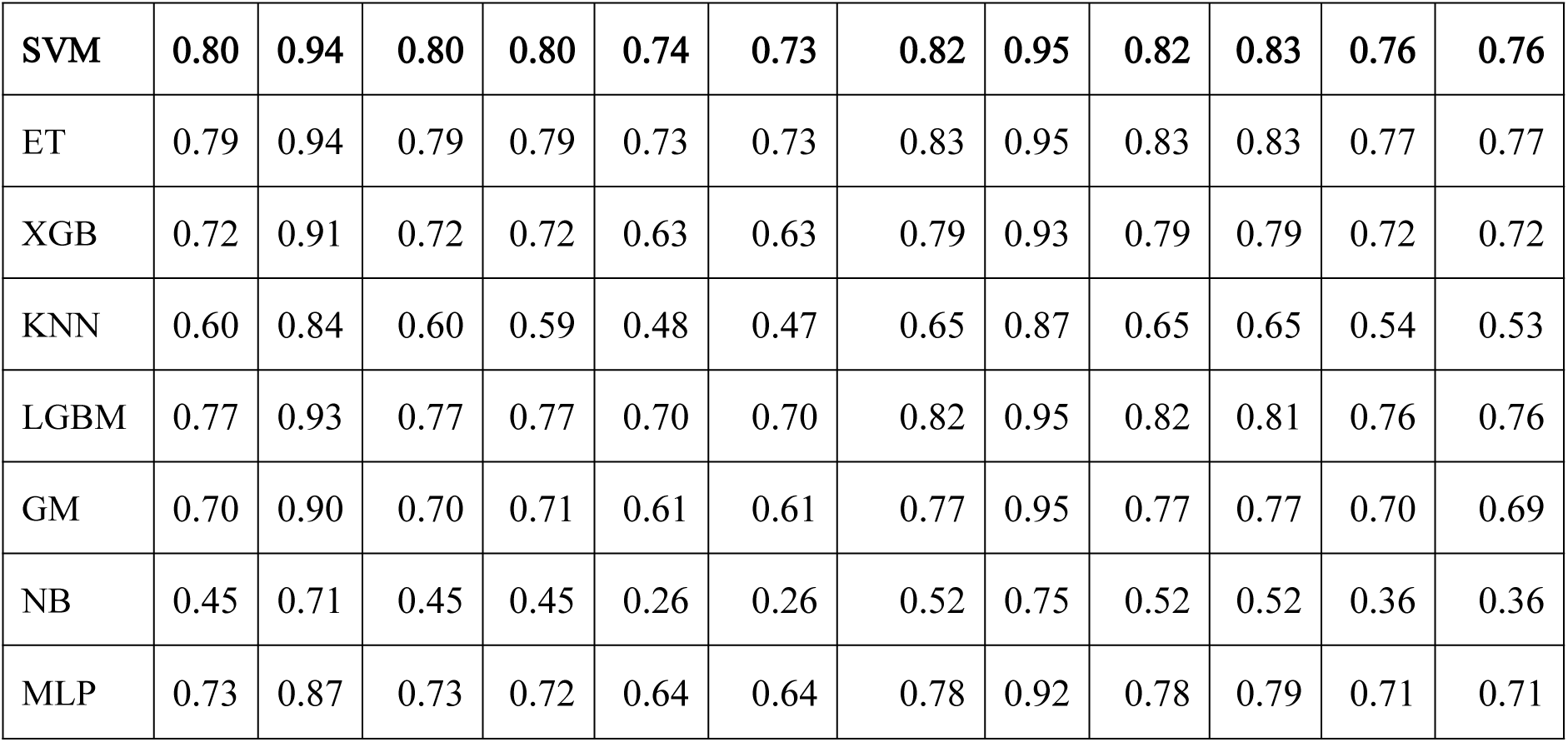
Per for mance on second Biomar ker Set with Clinical Features.

## Discussion

The incidence of thyroid cancer is rising, particularly among females, and while environmental factors are being explored, advancements in detection techniques likely contributed to this increase [32]. Given the complexity of thyroid cancer progression and prognosis, identifying reliable biomarkers is essential for improving patient outcomes. In this study, we analyzed transcriptomic data from 572 patients with available survival information using two approaches traditional statistical methods and machine learning-based techniques to identify distinct sets of prognostic biomarkers. We first conducted a correlation analysis, which revealed that MAFF and NR4A3 had positive correlations with overall survival, with correlation coefficients of 0.25 and 0.24, respectively, indicating that higher expression of these genes is associated with improved survival. Conversely, LOC728264 and VAMP1 exhibited negative correlations (-0.21 and -0.20), suggesting that their increased expression is linked to poorer outcomes. Subsequently, survival analysis identified 883 genes significantly associated with OS (p < 0.01), among which 541 had hazard ratios (HR) greater than 1, indicating potential risk factors, while 342 had HRs less than 1, suggesting protective roles. In lasso analysis we found 31 genes with non-zero coef value.Among the identified genes, 13 acted as protective factors with negative beta coefficient values, while 17 were associated with elevated risk, indicated by positive coefficient values. Elevated STC1 expression in thyroid lesions may serve as a useful diagnostic marker for papillary thyroid carcinoma (PTC) and is linked to lymph node metastasis risk, indicating its potential as a prognostic factor [33]. STC1 is also among genes highly upregulated in thyroid side population (SP) cells relative to non-SP cells, consistent with its role in tumorigenesis [34]. Overexpression of FOLR1 has been widely studied in breast, lung, endometrial, and ovarian cancers, where it correlates with cancer progression and reduced survival [35]. TFRC and TF are associated with poorer prognosis in thyroid cancer prediction models, with TFRC inducing ferroptosis and inhibiting tumor cell proliferation [36]. The gene MYL3 correlates with poor overall survival in Ewing sarcoma, and MYL9 is linked to aggressive glioblastoma, though no significant role in thyroid cancer (THCA) has been reported [37]. AKR1C3 is upregulated in thyroid cancer cells; its silencing induces cell cycle arrest and apoptosis in TPC-1 cells [38]. The long non-coding RNA KCNMB2-AS1 promotes ovarian cancer proliferation, migration, and inhibits apoptosis, suggesting its oncogenic role [39]. SLC19A1 is aberrantly expressed in multiple myeloma and is identified as a hypoxia-immune related gene linked to poor prognosis [40]. F12 shows high diagnostic efficacy in PTC and serves as an independent prognostic marker, involving metabolic pathways such as glutathione metabolism [41]. GRIN2A mutations are occasionally found in thyroid cancer cell lines [42], while Ubtor/KIAA1024 regulate mTOR signaling and cellular growth, representing potential therapeutic targets [43]. Overexpression of WT1 was observed in primary thyroid cancers [44]. Promoter hypomethylation of genes such as PCDHA5 in colorectal cancer, involved in cadherin pathways, hints at epigenetic regulation mechanisms potentially relevant to thyroid cancer [45]. The KIAA1644 gene is upregulated in thyroid cancer cells following metformin treatment. Beta IV tubulin isotypes TUBB4 and TUBB2C are highly expressed in pancreatic cancer cells and influence growth and chemotherapy response, but their role in THCA remains unknown. Members of the HtrA serine protease family show varied roles in cancers: HtrA1 and HtrA3 act as tumor suppressors, HtrA2 is tumor-type dependent, and HtrA4’s role is unclear [46]. ASCL2 is upregulated and hypomethylated in gastric cancer, correlating inversely with patient survival [47]. Overexpression of ZNF367 distinguishes PTC from benign and normal thyroid tissue [48]. RPS27 regulates expression and splicing of genes involved in inflammatory and toll-like receptor pathways linked to thyroid cancer progression [49].Moreover, we identified 233 drugs that target 13 of the genes highlighted in our analysis.GRIN2A, linked to poor prognosis, is inhibited by 69 drugs, while F12, MYL3, TFRC, and FOLR3 are targeted by 6, 3, 3, and 5 inhibitors respectively. These findings reveal promising druggable targets for developing therapies for high-risk thyroid cancer patients.

For the machine learning-based analysis, we applied feature selection techniques to identify key genes and obtained seven distinct sets of 20 prognostic biomarkers, each demonstrating strong predictive performance with favorable AUC values. Several primary biomarkers identified include ESRRB, LOC389705, GRM3, FLJ39582, C1orf168, PHYHIPL, PPIAL4D, ELOVL3, TMPRSS5, CXADRP3, SRMS, C7orf13, PCDHGB3, FAM153A, TH, LOC100133669, TMEM26, TCP11, STC1, and HSD17B14. Among these, GRM3 acts as an oncogene in glioma, kidney, and melanoma but has not been linked to THCA [50]. Notably, TMEM26 is identified as a rare fusion partner of RET in thyroid cancer, implicating it in tumorigenesis [51]. TCP11 is highly expressed in cervical cancer and correlates positively with patient survival but lacks data in thyroid cancer [52]. HSD17B14 is strongly coexpressed with APOE in thyroid cancer, suggesting involvement in lipid or hormone pathways [53]. Secondary biomarkers identified include GHDC, LOC100133161, TRIB1, CBLN3, LOC100134368, KBTBD10, PM20D1, ACOXL, THOC3, SLC5A10, DCAF12L2, SPDYE2, LOC100131496, C6orf164, CD302, KSR2, HEPHL1, CCDC89, LRRC8E, and POU2F1. The expression of TRIB1 is linked to thyroid cancer etiology [54]. PM20D1 is downregulated or silenced in hepatocellular carcinoma and acute myeloid leukemia [55,56]. The oncogenic role of THOC3 has been described across cancers and is proposed as a tumor driver in thyroid cancer [57]. LRRC8E is upregulated in THCA [58]. POU2F1 mediates the oncogenic effects of SOX12, promoting proliferation, migration, invasion, and epithelial-mesenchymal transition (EMT) in thyroid cancer [59]. Collectively, these findings highlight a broad set of prognostic biomarkers for thyroid cancer diagnosis, prognosis, and therapeutic targeting. Although we successfully conducted survival analysis and machine learning-based prognostic modeling using the TCGA-THCA dataset, we aimed to validate our findings using independent external datasets from the GEO database. However, an exhaustive search of available thyroid cancer datasets in GEO did not yield any suitable entries containing overall survival time or vital status information. Specifically, we examined datasets such as GSE48953 and GSE53157, which include limited clinical parameters like age, gender, stage, and mutational status, but lack survival outcome data. Other commonly referenced datasets, including GSE3467, GSE3678, GSE33630, and GSE53157, previously used in thyroid cancer studies, also do not contain survival information. Consequently, external validation of the prognostic biomarkers identified in our study could not be performed using GEO data.

This study is further limited by its reliance on in silico data without experimental validation, which may affect the biological relevance of the findings. Moreover, the absence of suitable external datasets with survival information underscores the need for future experimental and clinical studies to validate and confirm these results.

## Conclusion

This study identified multiple prognostic biomarkers associated with thyroid cancer survival, including both potential risk and protective genes. Using an integrative approach that combined statistical and machine learning methods, we developed seven robust gene sets with strong predictive power. Importantly, incorporating clinical features such as age, gender, and stage further improved model performance, demonstrating the added value of combining molecular and clinical data. These findings offer valuable insights into thyroid cancer biology and provide promising candidates for future diagnostic and therapeutic development.

## Authors’ Contributions

SM and GPSR collected the dataset. SM preprocessed the data. SM developed the classification models and implemented the algorithms. SM and GPSR analyzed the results. SM and GPSR drafted the manuscript. SM and GPSR reviewed and revised the manuscript. All authors have read and approved the final manuscript.

## Acknowledgments

The authors are grateful to the Department of Science and Technology (DST-INSPIRE) for providing fellowship and financial support, and to the Department of Computational Biology, IIIT Delhi, New Delhi, for providing infrastructure and facilities. The authors also acknowledge that the figures were created using BioRender.com, and that ChatGPT was used for language editing and refinement.

## Abbreviations

THCA: Thyroid Carcinoma
OS: Overall Survival
TCGA: The Cancer Genome Atlas
HR: Hazard Ratio
FDR: False Discovery Rate
DGIdb: Drug–Gene Interaction Database
ML: Machine Learning
AUROC: Area Under the Receiver Operating Characteristic Curve

