## Supplementary Tables (S1-S9), Supplementary Table S10 for "Integrative Bioinformatics Approach to Identify Prognostic Gene Signatures for Risk Stratification in Thyroid Carcinoma"

**Supplementary Table S1: The table shows the performance on 10 features selected using selectkbest approach.**

| Model | Train | | | | | | Test | | | | | |
| --- | --- | --- | --- | --- | --- | --- | --- | --- | --- | --- | --- | --- |
|  | **Acc** | **AUC** | **Sens** | **Spec** | **MCC** | **Kappa** | **Acc** | **AUC** | **Sens** | **Spec** | **MCC** | **Kappa** |
| RF | 0.62 | 0.84 | 0.62 | 0.62 | 0.49 | 0.49 | 0.63 | 0.83 | 0.63 | 0.63 | 0.51 | 0.50 |
| SVM | 0.61 | 0.83 | 0.61 | 0.60 | 0.49 | 0.49 | 0.64 | 0.82 | 0.64 | 0.64 | 0.53 | 0.52 |
| LR | 0.43 | 0.70 | 0.43 | 0.41 | 0.24 | 0.23 | 0.45 | 0.71 | 0.45 | 0.44 | 0.28 | 0.27 |
| ET | 0.67 | 0.87 | 0.67 | 0.67 | 0.57 | 0.56 | 0.66 | 0.86 | 0.66 | 0.65 | 0.54 | 0.54 |
| XGB | 0.61 | 0.82 | 0.61 | 0.60 | 0.48 | 0.47 | 0.58 | 0.82 | 0.58 | 0.57 | 0.45 | 0.44 |
| KNN | 0.56 | 0.80 | 0.56 | 0.56 | 0.42 | 0.41 | 0.53 | 0.80 | 0.53 | 0.53 | 0.39 | 0.38 |
| LGBM | 0.61 | 0.83 | 0.61 | 0.60 | 0.48 | 0.48 | 0.62 | 0.83 | 0.62 | 0.61 | 0.50 | 0.50 |
| GM | 0.62 | 0.84 | 0.62 | 0.62 | 0.50 | 0.49 | 0.58 | 0.83 | 0.58 | 0.57 | 0.44 | 0.44 |
| NB | 0.40 | 0.67 | 0.40 | 0.41 | 0.21 | 0.21 | 0.40 | 0.67 | 0.40 | 0.39 | 0.21 | 0.21 |
| MLP | 0.60 | 0.81 | 0.60 | 0.59 | 0.47 | 0.47 | 0.57 | 0.78 | 0.57 | 0.57 | 0.43 | 0.43 |

**Supplementary Table S2: The table shows the performance on 20 features selected using selectkbest approach.**

| Model | Train | | | | | | Test | | | | | |
| --- | --- | --- | --- | --- | --- | --- | --- | --- | --- | --- | --- | --- |
|  | **Acc** | **AUC** | **Sens** | **Spec** | **MCC** | **Kappa** | **Acc** | **AUC** | **Sens** | **Spec** | **MCC** | **Kappa** |
| RF | 0.65 | 0.87 | 0.65 | 0.65 | 0.54 | 0.54 | 0.70 | 0.86 | 0.70 | 0.69 | 0.60 | 0.60 |
| SVM | 0.73 | 0.89 | 0.73 | 0.73 | 0.65 | 0.64 | 0.72 | 0.88 | 0.72 | 0.71 | 0.62 | 0.62 |
| LR | 0.49 | 0.73 | 0.49 | 0.48 | 0.32 | 0.32 | 0.47 | 0.72 | 0.47 | 0.46 | 0.29 | 0.29 |
| ET | 0.71 | 0.90 | 0.71 | 0.71 | 0.62 | 0.62 | 0.73 | 0.89 | 0.73 | 0.73 | 0.65 | 0.65 |
| XGB | 0.65 | 0.86 | 0.65 | 0.65 | 0.54 | 0.54 | 0.68 | 0.86 | 0.68 | 0.68 | 0.57 | 0.57 |
| KNN | 0.59 | 0.81 | 0.59 | 0.59 | 0.46 | 0.45 | 0.57 | 0.82 | 0.57 | 0.54 | 0.44 | 0.43 |
| LGBM | 0.69 | 0.88 | 0.69 | 0.68 | 0.58 | 0.58 | 0.71 | 0.89 | 0.71 | 0.71 | 0.62 | 0.61 |
| GM | 0.65 | 0.87 | 0.65 | 0.65 | 0.54 | 0.54 | 0.70 | 0.88 | 0.70 | 0.69 | 0.60 | 0.60 |
| NB | 0.42 | 0.69 | 0.42 | 0.42 | 0.23 | 0.23 | 0.44 | 0.68 | 0.44 | 0.44 | 0.25 | 0.25 |
| MLP | 0.70 | 0.86 | 0.70 | 0.69 | 0.60 | 0.60 | 0.65 | 0.82 | 0.65 | 0.65 | 0.54 | 0.54 |

**Supplementary Table S3: The table shows the performance on 50 features selected using selectkbest approach.**

| Model | Train | | | | | | Test | | | | | |
| --- | --- | --- | --- | --- | --- | --- | --- | --- | --- | --- | --- | --- |
|  | **Acc** | **AUC** | **Sens** | **Spec** | **MCC** | **Kappa** | **Acc** | **AUC** | **Sens** | **Spec** | **MCC** | **Kappa** |
| RF | 0.72 | 0.90 | 0.72 | 0.71 | 0.62 | 0.62 | 0.70 | 0.89 | 0.70 | 0.70 | 0.60 | 0.60 |
| SVM | 0.77 | 0.94 | 0.77 | 0.77 | 0.70 | 0.70 | 0.79 | 0.93 | 0.79 | 0.79 | 0.72 | 0.72 |
| LR | 0.55 | 0.77 | 0.55 | 0.54 | 0.40 | 0.40 | 0.54 | 0.76 | 0.54 | 0.54 | 0.38 | 0.38 |
| ET | 0.75 | 0.92 | 0.75 | 0.75 | 0.67 | 0.66 | 0.73 | 0.91 | 0.73 | 0.75 | 0.65 | 0.65 |
| XGB | 0.71 | 0.90 | 0.71 | 0.71 | 0.62 | 0.62 | 0.68 | 0.88 | 0.68 | 0.68 | 0.58 | 0.58 |
| KNN | 0.59 | 0.83 | 0.59 | 0.58 | 0.47 | 0.46 | 0.61 | 0.84 | 0.61 | 0.60 | 0.49 | 0.47 |
| LGBM | 0.74 | 0.92 | 0.74 | 0.74 | 0.66 | 0.66 | 0.72 | 0.91 | 0.72 | 0.71 | 0.62 | 0.62 |
| GM | 0.70 | 0.90 | 0.70 | 0.69 | 0.60 | 0.60 | 0.71 | 0.90 | 0.71 | 0.70 | 0.61 | 0.61 |
| NB | 0.43 | 0.70 | 0.43 | 0.42 | 0.24 | 0.24 | 0.42 | 0.69 | 0.42 | 0.43 | 0.23 | 0.23 |
| MLP | 0.72 | 0.89 | 0.72 | 0.71 | 0.63 | 0.63 | 0.72 | 0.87 | 0.72 | 0.72 | 0.64 | 0.63 |

**Supplementary Table S4: The table shows the performance on 100 features selected using selectkbest approach.**

| Model | Train | | | | | | Test | | | | | |
| --- | --- | --- | --- | --- | --- | --- | --- | --- | --- | --- | --- | --- |
|  | **Acc** | **AUC** | **Sens** | **Spec** | **MCC** | **Kappa** | **Acc** | **AUC** | **Sens** | **Spec** | **MCC** | **Kappa** |
| RF | 0.73 | 0.91 | 0.73 | 0.74 | 0.65 | 0.65 | 0.74 | 0.91 | 0.74 | 0.76 | 0.66 | 0.66 |
| SVM | 0.81 | 0.96 | 0.81 | 0.81 | 0.75 | 0.75 | 0.83 | 0.94 | 0.83 | 0.83 | 0.78 | 0.78 |
| LR | 0.64 | 0.83 | 0.64 | 0.64 | 0.53 | 0.53 | 0.59 | 0.81 | 0.59 | 0.60 | 0.46 | 0.46 |
| ET | 0.76 | 0.93 | 0.76 | 0.77 | 0.69 | 0.69 | 0.77 | 0.93 | 0.77 | 0.78 | 0.70 | 0.69 |
| XGB | 0.74 | 0.92 | 0.74 | 0.74 | 0.66 | 0.65 | 0.76 | 0.92 | 0.76 | 0.76 | 0.68 | 0.68 |
| KNN | 0.61 | 0.84 | 0.61 | 0.61 | 0.50 | 0.48 | 0.62 | 0.85 | 0.62 | 0.60 | 0.51 | 0.50 |
| LGBM | 0.77 | 0.94 | 0.77 | 0.77 | 0.70 | 0.70 | 0.78 | 0.93 | 0.78 | 0.78 | 0.71 | 0.71 |
| GM | 0.68 | 0.87 | 0.68 | 0.68 | 0.58 | 0.58 | 0.71 | 0.90 | 0.71 | 0.71 | 0.61 | 0.61 |
| NB | 0.44 | 0.71 | 0.44 | 0.44 | 0.26 | 0.25 | 0.45 | 0.70 | 0.45 | 0.45 | 0.27 | 0.27 |
| MLP | 0.79 | 0.92 | 0.79 | 0.79 | 0.72 | 0.71 | 0.76 | 0.90 | 0.76 | 0.76 | 0.68 | 0.68 |

**Supplementary Table S5: The table shows the performance on 10 features selected using RFE method**

| Model | Train | | | | | | Test | | | | | |
| --- | --- | --- | --- | --- | --- | --- | --- | --- | --- | --- | --- | --- |
|  | **Acc** | **AUC** | **Sens** | **Spec** | **MCC** | **Kappa** | **Acc** | **AUC** | **Sens** | **Spec** | **MCC** | **Kappa** |
| RF | 0.74 | 0.91 | 0.74 | 0.74 | 0.66 | 0.66 | 0.72 | 0.89 | 0.72 | 0.71 | 0.63 | 0.63 |
| SVM | 0.77 | 0.93 | 0.77 | 0.77 | 0.70 | 0.70 | 0.79 | 0.93 | 0.79 | 0.79 | 0.73 | 0.72 |
| LR | 0.63 | 0.83 | 0.63 | 0.62 | 0.50 | 0.50 | 0.53 | 0.76 | 0.53 | 0.51 | 0.38 | 0.38 |
| ET | 0.77 | 0.93 | 0.77 | 0.77 | 0.69 | 0.69 | 0.75 | 0.92 | 0.75 | 0.75 | 0.67 | 0.67 |
| XGB | 0.71 | 0.90 | 0.71 | 0.71 | 0.62 | 0.62 | 0.67 | 0.88 | 0.67 | 0.65 | 0.56 | 0.56 |
| KNN | 0.62 | 0.85 | 0.62 | 0.61 | 0.50 | 0.49 | 0.66 | 0.86 | 0.66 | 0.63 | 0.55 | 0.54 |
| LGBM | 0.74 | 0.92 | 0.74 | 0.74 | 0.66 | 0.66 | 0.71 | 0.90 | 0.71 | 0.70 | 0.62 | 0.61 |
| GM | 0.68 | 0.88 | 0.68 | 0.68 | 0.57 | 0.57 | 0.68 | 0.89 | 0.68 | 0.67 | 0.58 | 0.58 |
| NB | 0.52 | 0.75 | 0.52 | 0.51 | 0.37 | 0.36 | 0.50 | 0.74 | 0.50 | 0.47 | 0.33 | 0.33 |
| MLP | 0.73 | 0.87 | 0.73 | 0.73 | 0.64 | 0.64 | 0.72 | 0.86 | 0.72 | 0.72 | 0.64 | 0.63 |

**Supplementary Table S6: The table shows the performance on 20 features selected using RFE method**

| Model | Train | | | | | | Test | | | | | |
| --- | --- | --- | --- | --- | --- | --- | --- | --- | --- | --- | --- | --- |
|  | **Acc** | **AUC** | **Sens** | **Spec** | **MCC** | **Kappa** | **Acc** | **AUC** | **Sens** | **Spec** | **MCC** | **Kappa** |
| RF | 0.74 | 0.91 | 0.74 | 0.74 | 0.66 | 0.66 | 0.72 | 0.89 | 0.72 | 0.71 | 0.63 | 0.63 |
| SVM | 0.77 | 0.93 | 0.77 | 0.77 | 0.70 | 0.70 | 0.79 | 0.93 | 0.79 | 0.79 | 0.73 | 0.72 |
| LR | 0.63 | 0.83 | 0.63 | 0.62 | 0.50 | 0.50 | 0.53 | 0.76 | 0.53 | 0.51 | 0.38 | 0.38 |
| ET | 0.77 | 0.93 | 0.77 | 0.77 | 0.69 | 0.69 | 0.75 | 0.92 | 0.75 | 0.75 | 0.67 | 0.67 |
| XGB | 0.71 | 0.90 | 0.71 | 0.71 | 0.62 | 0.62 | 0.67 | 0.88 | 0.67 | 0.65 | 0.56 | 0.56 |
| KNN | 0.62 | 0.85 | 0.62 | 0.61 | 0.50 | 0.49 | 0.66 | 0.86 | 0.66 | 0.63 | 0.55 | 0.54 |
| LGBM | 0.74 | 0.92 | 0.74 | 0.74 | 0.66 | 0.66 | 0.71 | 0.90 | 0.71 | 0.70 | 0.62 | 0.61 |
| GM | 0.68 | 0.88 | 0.68 | 0.68 | 0.57 | 0.57 | 0.68 | 0.89 | 0.68 | 0.67 | 0.58 | 0.58 |
| NB | 0.52 | 0.75 | 0.52 | 0.51 | 0.37 | 0.36 | 0.50 | 0.74 | 0.50 | 0.47 | 0.33 | 0.33 |
| MLP | 0.73 | 0.87 | 0.73 | 0.73 | 0.64 | 0.64 | 0.72 | 0.86 | 0.72 | 0.72 | 0.64 | 0.63 |

**Supplementary Table S7: The table shows the performance on 10 features selected using SVC-L1 method**

| Model | Train | | | | | | Test | | | | | |
| --- | --- | --- | --- | --- | --- | --- | --- | --- | --- | --- | --- | --- |
|  | **Acc** | **AUC** | **Sens** | **Spec** | **MCC** | **Kappa** | **Acc** | **AUC** | **Sens** | **Spec** | **MCC** | **Kappa** |
| RF | 0.64 | 0.86 | 0.64 | 0.65 | 0.53 | 0.53 | 0.61 | 0.84 | 0.61 | 0.62 | 0.49 | 0.49 |
| SVM | 0.66 | 0.86 | 0.66 | 0.66 | 0.55 | 0.55 | 0.65 | 0.84 | 0.65 | 0.64 | 0.53 | 0.53 |
| LR | 0.40 | 0.67 | 0.40 | 0.39 | 0.20 | 0.19 | 0.39 | 0.67 | 0.39 | 0.39 | 0.19 | 0.19 |
| ET | 0.69 | 0.89 | 0.69 | 0.69 | 0.59 | 0.58 | 0.69 | 0.87 | 0.69 | 0.69 | 0.58 | 0.58 |
| XGB | 0.64 | 0.85 | 0.64 | 0.64 | 0.52 | 0.52 | 0.63 | 0.84 | 0.63 | 0.63 | 0.51 | 0.50 |
| KNN | 0.56 | 0.80 | 0.56 | 0.57 | 0.43 | 0.41 | 0.56 | 0.80 | 0.56 | 0.55 | 0.43 | 0.42 |
| LGBM | 0.65 | 0.86 | 0.65 | 0.65 | 0.53 | 0.53 | 0.64 | 0.85 | 0.64 | 0.64 | 0.52 | 0.52 |
| GM | 0.64 | 0.85 | 0.64 | 0.64 | 0.52 | 0.52 | 0.64 | 0.85 | 0.64 | 0.64 | 0.52 | 0.52 |
| NB | 0.41 | 0.67 | 0.41 | 0.42 | 0.21 | 0.21 | 0.42 | 0.68 | 0.42 | 0.43 | 0.23 | 0.23 |
| MLP | 0.59 | 0.80 | 0.59 | 0.59 | 0.46 | 0.45 | 0.64 | 0.79 | 0.64 | 0.63 | 0.52 | 0.52 |

**Supplementary Table S8: The table shows the performance on 50 features selected using SVC-L1 method**

| Model | Train | | | | | | Test | | | | | |
| --- | --- | --- | --- | --- | --- | --- | --- | --- | --- | --- | --- | --- |
|  | **Acc** | **AUC** | **Sens** | **Spec** | **MCC** | **Kappa** | **Acc** | **AUC** | **Sens** | **Spec** | **MCC** | **Kappa** |
| RF | 0.81 | 0.95 | 0.81 | 0.81 | 0.75 | 0.75 | 0.78 | 0.94 | 0.78 | 0.78 | 0.71 | 0.71 |
| SVM | 0.88 | 0.98 | 0.88 | 0.88 | 0.84 | 0.84 | 0.83 | 0.96 | 0.83 | 0.83 | 0.77 | 0.77 |
| LR | 0.74 | 0.89 | 0.74 | 0.74 | 0.65 | 0.65 | 0.67 | 0.83 | 0.67 | 0.67 | 0.56 | 0.56 |
| ET | 0.84 | 0.96 | 0.84 | 0.85 | 0.79 | 0.79 | 0.81 | 0.95 | 0.81 | 0.81 | 0.75 | 0.75 |
| XGB | 0.82 | 0.95 | 0.82 | 0.82 | 0.76 | 0.75 | 0.79 | 0.94 | 0.79 | 0.79 | 0.72 | 0.72 |
| KNN | 0.64 | 0.87 | 0.64 | 0.65 | 0.55 | 0.52 | 0.67 | 0.88 | 0.67 | 0.70 | 0.59 | 0.57 |
| LGBM | 0.82 | 0.96 | 0.82 | 0.82 | 0.76 | 0.76 | 0.80 | 0.95 | 0.80 | 0.79 | 0.73 | 0.73 |
| GM | 0.66 | 0.87 | 0.66 | 0.66 | 0.55 | 0.55 | 0.74 | 0.91 | 0.74 | 0.74 | 0.66 | 0.65 |
| NB | 0.57 | 0.81 | 0.57 | 0.58 | 0.43 | 0.43 | 0.59 | 0.81 | 0.59 | 0.58 | 0.46 | 0.46 |
| MLP | 0.84 | 0.95 | 0.84 | 0.85 | 0.80 | 0.79 | 0.78 | 0.90 | 0.78 | 0.78 | 0.71 | 0.71 |

**Supplementary Table S9: The table shows the performance on 100 features selected using SVC-L1 method**

| Model | Train | | | | | | Test | | | | | |
| --- | --- | --- | --- | --- | --- | --- | --- | --- | --- | --- | --- | --- |
|  | **Acc** | **AUC** | **Sens** | **Spec** | **MCC** | **Kappa** | **Acc** | **AUC** | **Sens** | **Spec** | **MCC** | **Kappa** |
| RF | 0.83 | 0.96 | 0.83 | 0.84 | 0.78 | 0.78 | 0.81 | 0.95 | 0.81 | 0.81 | 0.74 | 0.74 |
| SVM | 0.91 | 0.99 | 0.91 | 0.91 | 0.88 | 0.88 | 0.84 | 0.97 | 0.84 | 0.84 | 0.79 | 0.79 |
| LR | 0.85 | 0.95 | 0.85 | 0.86 | 0.80 | 0.80 | 0.79 | 0.89 | 0.79 | 0.79 | 0.73 | 0.72 |
| ET | 0.84 | 0.97 | 0.84 | 0.84 | 0.78 | 0.78 | 0.83 | 0.96 | 0.83 | 0.83 | 0.77 | 0.77 |
| XGB | 0.81 | 0.96 | 0.81 | 0.81 | 0.74 | 0.74 | 0.78 | 0.94 | 0.78 | 0.78 | 0.71 | 0.71 |
| KNN | 0.63 | 0.85 | 0.63 | 0.68 | 0.53 | 0.51 | 0.69 | 0.87 | 0.69 | 0.72 | 0.62 | 0.59 |
| LGBM | 0.82 | 0.97 | 0.82 | 0.83 | 0.77 | 0.77 | 0.85 | 0.96 | 0.85 | 0.85 | 0.80 | 0.80 |
| GM | 0.68 | 0.88 | 0.68 | 0.68 | 0.58 | 0.58 | 0.74 | 0.91 | 0.74 | 0.74 | 0.66 | 0.66 |
| NB | 0.61 | 0.83 | 0.61 | 0.61 | 0.48 | 0.48 | 0.62 | 0.83 | 0.62 | 0.61 | 0.49 | 0.49 |
| MLP | 0.88 | 0.97 | 0.88 | 0.89 | 0.85 | 0.84 | 0.83 | 0.93 | 0.83 | 0.83 | 0.78 | 0.77 |

**Supplementary Table S10: The table shows the lists of drugs that are found to targets genes**

| **Gene** | **Drug** | **Regulatory approval** | **Indication** | **Interaction score** |
| --- | --- | --- | --- | --- |
| TFRC | E2.3 | Not Approved |  | 8.700633093 |
| TFRC | PABINAFUSP ALFA | Not Approved |  | 8.700633093 |
| TFRC | A27.15 | Not Approved |  | 17.40126619 |
| PZP | 4-PHENYLENEDIAMINE | Not Approved |  | 2.485895169 |
| PZP | PROGESTERONE | Approved | for reducing the risk of pre-term birth for women with short cervix a mid-pregnancy,for prevention of preterm delivery,for symptomatic treatment of menopausal symptoms,neuroprotectant for stroke victims | 0.294936715 |
| PZP | ESTRADIOL VALERATE | Approved | for treatment of menopausal symptoms,contraceptive,treatment for menopause,hormone replacement | 0.370239706 |
| PZP | ESTRAMUSTINE PHOSPHATE | Not Approved |  | 8.700633093 |
| PZP | NORETHINDRONE | Approved | Synthetic,Oral,Contraceptives,contraceptive | 1.740126619 |
| PZP | DEXAMETHASONE | Approved | for treatment of Meniere's disease,glucocorticoid,antiinflammatory agent | 0.204720779 |
| IL5RA | BENRALIZUMAB | Approved | antiasthmatic agent | 52.20379856 |
| RPS27 | DORLIMOMAB ARITOX | Not Approved |  | 0.093221069 |
| RPS27 | CYCLOHEXIMIDE | Not Approved |  | 0.078501953 |
| RPS27 | ATALUREN | Approved |  | 0.090947384 |
| RPS27 | MT-3724 | Not Approved |  | 0.092070191 |
| RPS27 | EXALUREN | Not Approved |  | 0.093221069 |
| RPS27 | BOS172722 | Not Approved |  | 1.242947585 |
| RPS27 | EMPESERTIB | Not Approved |  | 2.485895169 |
| MYL3 | DANICAMTIV | Not Approved |  | 2.485895169 |
| MYL3 | MAVACAMTEN | Approved |  | 1.933474021 |
| MYL3 | OMECAMTIV MECARBIL | Not Approved | for treatment of heart failure | 2.175158273 |
| AKR1C3 | EXEMESTANE | Approved | antineoplastic agent | 0.435031655 |
| AKR1C3 | BACCHARIN | Not Approved |  | 6.960506475 |
| AKR1C3 | DAUNORUBICIN LIPOSOMAL | Approved | antineoplastic agent | 0.076489082 |
| AKR1C3 | INDOMETHACIN | Approved | NSAID | 0.174012662 |
| AKR1C3 | ANTI-ANDROGEN | Not Approved |  | 1.740126619 |
| AKR1C3 | NONSTEROIDAL ANTIINFLAMMATORY DRUG | Not Approved |  | 0.614162336 |
| AKR1C3 | DOXORUBICIN HYDROCHLORIDE | Approved | antineoplastic agent | 0.070545674 |
| AKR1C3 | CHEMBL:CHEMBL1682201 | Not Approved |  | 6.960506475 |
| AKR1C3 | DOCETAXEL ANHYDROUS | Approved | antineoplastic agent | 0.083861524 |
| AKR1C3 | 17BETA-HYDROXYSTEROID DEHYDROGENASE TYPE 5 INHIBITOR ASP9521 | Not Approved |  | 3.480253237 |
| AKR1C3 | CHEMBL:CHEMBL1682200 | Not Approved |  | 6.960506475 |
| AKR1C3 | CHEMBL:CHEMBL1682202 | Not Approved |  | 6.960506475 |
| AKR1C3 | TESTOSTERONE | Approved | for treatment of female sexual dysfunction,hormone replacement | 0.178474525 |
| AKR1C3 | THERAPEUTIC ANDROGEN | Not Approved |  | 0.409441557 |
| AKR1C3 | FLUFENAMIC ACID | Not Approved |  | 0.091585612 |
| KCNMB2 | RITODRINE | Approved | Tocolytic Agents | 34.80253237 |
| SLC19A1 | CAPECITABINE | Approved |  | 0.113733766 |
| SLC19A1 | IMATINIB | Approved | antineoplastic agent | 0.080561418 |
| SLC19A1 | RALTITREXED | Approved | antineoplastic agent | 1.657263446 |
| SLC19A1 | PEMETREXED DISODIUM | Approved | Antineoplastic Agents,antineoplastic agent | 0.669279469 |
| SLC19A1 | CYCLOSPORINE | Approved | immunosuppressant,opthalmological agent | 0.066671518 |
| SLC19A1 | FOLIC ACID | Approved |  | 0.362526379 |
| SLC19A1 | CISPLATIN | Approved |  | 0.039458653 |
| SLC19A1 | LOMETREXOL | Not Approved |  | 1.160084412 |
| SLC19A1 | PREDNISONE | Approved | corticosteroid,antiinflammatory agent | 0.128898268 |
| SLC19A1 | SULFASALAZINE | Approved | DMARD,antiinflammatory agent | 0.241684253 |
| SLC19A1 | METHOTREXATE | Approved | DMARD | 0.026365555 |
| SLC19A1 | CARBOPLATIN | Approved |  | 0.067446768 |
| SLC19A1 | TALOTREXIN | Not Approved | antineoplastic agent | 1.933474021 |
| SLC19A1 | BEVACIZUMAB | Approved | antineoplastic agent | 0.145010552 |
| SLC19A1 | HYDROXYCHLOROQUINE | Approved | antirheumatic agent | 0.32224567 |
| SLC19A1 | IRINOTECAN HYDROCHLORIDE | Approved | antineoplastic agent | 0.092070191 |
| SLC19A1 | FLUOROURACIL | Approved |  | 0.075005458 |
| SLC19A1 | LEUCOVORIN CALCIUM | Approved | adjuvant to chemotherapy | 0.483368505 |
| F12 | MTK458 | Not Approved |  | 8.700633093 |
| F12 | PENTAMIDINE ISETHIONATE | Approved |  | 1.242947585 |
| F12 | BISMUTH SUBGALLATE | Approved |  | 8.700633093 |
| F12 | BORTEZOMIB | Approved | antineoplastic agent | 0.204720779 |
| F12 | HUMAN C1-ESTERASE INHIBITOR | Approved |  | 2.175158273 |
| F12 | ANTITHROMBIN ALFA | Approved | antithrombotic | 0.96673701 |
| GRIN2A | ORPHENADRINE CITRATE | Approved |  | 0.036027466 |
| GRIN2A | PHILANTHOTOXIN 343 | Not Approved |  | 0.504384527 |
| GRIN2A | AZD8108 | Not Approved |  | 0.108082399 |
| GRIN2A | RALFINAMIDE | Not Approved |  | 0.03782884 |
| GRIN2A | CONANTOKIN-G | Not Approved |  | 0.189144198 |
| GRIN2A | NVP-AAM077 | Not Approved |  | 0.378288395 |
| GRIN2A | GW468816 | Not Approved |  | 0.108082399 |
| GRIN2A | APIMOSTINEL | Not Approved |  | 0.108082399 |
| GRIN2A | SELFOTEL | Not Approved |  | 0.189144198 |
| GRIN2A | MEMANTINE HYDROCHLORIDE | Approved | for treatment of Alzheimer's disease,for treatment of glaucoma | 0.420320439 |
| GRIN2A | FELBAMATE | Approved | Anticonvulsants; Antiepileptics | 0.189144198 |
| GRIN2A | NEBOGLAMINE | Not Approved |  | 0.108082399 |
| GRIN2A | ZOLBETUXIMAB | Not Approved |  | 0.756576791 |
| GRIN2A | GV196771A | Not Approved |  | 0.189144198 |
| GRIN2A | [3H]CGP61594 | Not Approved |  | 0.151315358 |
| GRIN2A | [3H]DIZOCILPINE | Not Approved |  | 0.151315358 |
| GRIN2A | (RS)-(TETRAZOL-5-YL)GLYCINE | Not Approved |  | 0.151315358 |
| GRIN2A | L689560 | Not Approved |  | 0.151315358 |
| GRIN2A | PERZINFOTEL | Not Approved |  | 0.108082399 |
| GRIN2A | AV-101 | Not Approved |  | 0.108082399 |
| GRIN2A | GLYCINE | Approved |  | 0.126096132 |
| GRIN2A | KETAMINE HYDROCHLORIDE | Approved | analgesic | 0.178018068 |
| GRIN2A | MG2+ | Not Approved |  | 0.058198215 |
| GRIN2A | NERAMEXANE | Not Approved | for treatment of Alzheimer's disease | 0.252192264 |
| GRIN2A | D-AP5 | Not Approved |  | 0.189144198 |
| GRIN2A | NBQX | Not Approved |  | 0.047286049 |
| GRIN2A | L701324 | Not Approved |  | 0.126096132 |
| GRIN2A | LANICEMINE | Not Approved | antidepressant | 0.216164797 |
| GRIN2A | NERAMEXANE MESYLATE | Not Approved |  | 0.084064088 |
| GRIN2A | CGP37849 | Not Approved |  | 0.189144198 |
| GRIN2A | ESKETAMINE | Approved |  | 0.108082399 |
| GRIN2A | INDANTADOL | Not Approved | neuropathic pain,analgesic | 0.084064088 |
| GRIN2A | D-CCPENE | Not Approved |  | 0.189144198 |
| GRIN2A | DEXANABINOL | Not Approved | neuroprotectant | 0.126096132 |
| GRIN2A | [3H]D-ASPARTIC ACID | Not Approved |  | 0.068779708 |
| GRIN2A | TCN-201 | Not Approved |  | 0.756576791 |
| GRIN2A | LY233053 | Not Approved |  | 0.189144198 |
| GRIN2A | SAGE-718 | Not Approved |  | 0.378288395 |
| GRIN2A | (+)-HA966 | Not Approved |  | 0.252192264 |
| GRIN2A | NMDA | Not Approved |  | 0.084064088 |
| GRIN2A | CNS-5161 | Not Approved | analgesic | 0.216164797 |
| GRIN2A | AMANTADINE HYDROCHLORIDE | Approved |  | 0.151315358 |
| GRIN2A | DIMIRACETAM | Not Approved | nootropic | 0.189144198 |
| GRIN2A | INDANTADOL | Not Approved |  | 0.094572099 |
| GRIN2A | NYX-2925 | Not Approved |  | 0.756576791 |
| GRIN2A | 5,7-DICHLOROKYNURENIC ACID | Not Approved |  | 0.108082399 |
| GRIN2A | DELUCEMINE | Not Approved |  | 0.108082399 |
| GRIN2A | ASPARTIC ACID | Approved |  | 0.075657679 |
| GRIN2A | DEXTROMETHORPHAN HYDROBROMIDE | Approved |  | 0.378288395 |
| GRIN2A | [3H]MDL105519 | Not Approved |  | 0.189144198 |
| GRIN2A | DEXTROMETHORPHAN POLISTIREX | Approved | Antitussive Agents,analgesic,antitussive agent | 0.047286049 |
| GRIN2A | [3H]CGS19755 | Not Approved |  | 0.151315358 |
| GRIN2A | [3H]L689560 | Not Approved |  | 0.151315358 |
| GRIN2A | UBP141 | Not Approved |  | 0.189144198 |
| GRIN2A | ACAMPROSATE CALCIUM | Approved | antineoplastic agent,for treatment of alcohol-dependance | 0.030263072 |
| GRIN2A | ORPHENADRINE HYDROCHLORIDE | Approved |  | 0.039819831 |
| GRIN2A | ZWB67 | Not Approved |  | 0.378288395 |
| GRIN2A | PHENCYCLIDINE | Not Approved |  | 0.189144198 |
| GRIN2A | DIZOCILPINE | Not Approved |  | 0.378288395 |
| GRIN2A | ESKETAMINE HYDROCHLORIDE | Not Approved | for treatment of tinnitus | 0.108082399 |
| GRIN2A | LATREPIRDINE | Not Approved | neuroprotectant | 0.094572099 |
| GRIN2A | [3H]CGP39653 | Not Approved |  | 0.151315358 |
| GRIN2A | N1-DANSYL-SPERMINE | Not Approved |  | 0.189144198 |
| GRIN2A | RAPASTINEL | Not Approved |  | 0.108082399 |
| GRIN2A | HOMOQUINOLINIC ACID | Not Approved |  | 0.151315358 |
| GRIN2A | SERINE, D- | Not Approved |  | 0.126096132 |
| GRIN2A | HALOPERIDOL DECANOATE | Approved | Antipsychotic Agents | 0.019909916 |
| GRIN2A | PLAZINEMDOR | Not Approved |  | 0.756576791 |
| GRIN2A | [3H]CPP | Not Approved |  | 0.151315358 |
| WT1 | CURCUMIN | Approved |  | 0.121687176 |
| WT1 | GALINPEPIMUT-S | Not Approved |  | 4.015676812 |
| WT1 | TRETINOIN | Approved | for treatment of acne | 0.13238495 |
| WT1 | HALOFUGINONE | Not Approved | antineoplastic agent | 1.338558937 |
| WT1 | RECOMBINANT CYTOKINE | Not Approved |  | 0.308898216 |
| WT1 | CYTARABINE | Approved | antineoplastic agent | 0.05019596 |
| WT1 | SIROLIMUS | Approved | for treatment of wet age-related macular degeneration,immunosuppressant | 0.138471614 |
| WT1 | DAUNORUBICIN LIPOSOMAL | Approved | antineoplastic agent | 0.044128317 |
| WT1 | DIMETHYL SULFOXIDE | Approved |  | 0.501959602 |
| WT1 | RECOMBINANT 70-KD HEAT-SHOCK PROTEIN | Not Approved |  | 0.803135362 |
| WT1 | IH636 GRAPE SEED PROANTHOCYANIDIN EXTRACT | Not Approved |  | 1.606270725 |
| WT1 | DEFEROXAMINE | Approved |  | 0.472432566 |
| WT1 | OMBIPEPIMUT-S | Not Approved |  | 4.015676812 |
| TUBB4B | SOFITUZUMAB VEDOTIN | Not Approved |  | 0.038842112 |
| TUBB4B | CHEMBL:CHEMBL2036124 | Not Approved |  | 0.077684224 |
| TUBB4B | VORINOSTAT | Approved | antineoplastic agent | 0.022195493 |
| TUBB4B | DOLASTATIN-10 | Not Approved |  | 0.038842112 |
| TUBB4B | CHEMBL:CHEMBL1935538 | Not Approved |  | 0.073114564 |
| TUBB4B | ABT-751 | Not Approved |  | 0.109671846 |
| TUBB4B | SAGOPILONE | Not Approved |  | 0.041431586 |
| TUBB4B | VINCRISTINE SULFATE | Approved |  | 0.032709147 |
| TUBB4B | ASG-5ME | Not Approved |  | 0.038842112 |
| TUBB4B | VANDORTUZUMAB VEDOTIN | Not Approved |  | 0.038842112 |
| TUBB4B | LORVOTUZUMAB MERTANSINE | Not Approved |  | 0.038842112 |
| TUBB4B | CABAZITAXEL | Approved | antineoplastic agent | 0.028248809 |
| TUBB4B | TISOTUMAB VEDOTIN | Approved |  | 0.038842112 |
| TUBB4B | INDUSATUMAB VEDOTIN | Not Approved |  | 0.038842112 |
| TUBB4B | AZINTUXIZUMAB VEDOTIN | Not Approved |  | 0.038842112 |
| TUBB4B | DOCETAXEL ANHYDROUS | Approved | antineoplastic agent | 0.007487636 |
| TUBB4B | AGS-16C3F | Not Approved |  | 0.038842112 |
| TUBB4B | ZAMPANOLIDE | Not Approved |  | 0.077684224 |
| TUBB4B | PODOFILOX | Approved | Phytogenic; Keratolytic Agents,Antineoplastic Agents | 0.103578965 |
| TUBB4B | CHEMBL:CHEMBL2036119 | Not Approved |  | 0.077684224 |
| TUBB4B | PLOCABULIN | Not Approved |  | 0.041431586 |
| TUBB4B | FOSBRETABULIN DISODIUM | Not Approved |  | 0.036557282 |
| TUBB4B | ENAPOTAMAB VEDOTIN | Not Approved |  | 0.038842112 |
| TUBB4B | SOBLIDOTIN | Not Approved |  | 0.036557282 |
| TUBB4B | VINFLUNINE DITARTRATE | Approved |  | 0.041431586 |
| TUBB4B | CHEMBL:CHEMBL1795737 | Not Approved |  | 0.077684224 |
| TUBB4B | OMBRABULIN | Not Approved | antineoplastic agent | 0.041431586 |
| TUBB4B | TELISOTUZUMAB VEDOTIN | Not Approved |  | 0.038842112 |
| TUBB4B | PACLITAXEL DOCOSAHEXAENOIC ACID | Not Approved |  | 0.041431586 |
| TUBB4B | BIVATUZUMAB MERTANSINE | Not Approved |  | 0.038842112 |
| TUBB4B | VINORELBINE TARTRATE | Approved |  | 0.041431586 |
| TUBB4B | RG-7841 | Not Approved |  | 0.038842112 |
| TUBB4B | COLCHICINE | Approved | for treatment of gout | 0.107562772 |
| TUBB4B | RG-7636 | Not Approved |  | 0.038842112 |
| TUBB4B | NOCODAZOLE | Not Approved |  | 0.155368448 |
| TUBB4B | IXABEPILONE | Approved | antineoplastic agent | 0.028248809 |
| TUBB4B | ENFORTUMAB VEDOTIN | Approved |  | 0.036557282 |
| TUBB4B | LARGAZOLE | Not Approved |  | 0.042860262 |
| TUBB4B | BMS-275183 | Not Approved |  | 0.038842112 |
| TUBB4B | MIRVETUXIMAB SORAVTANSINE | Approved |  | 0.038842112 |
| TUBB4B | ERIBULIN MESYLATE | Approved |  | 0.025894741 |
| TUBB4B | CHEMBL:CHEMBL453818 | Not Approved |  | 0.077684224 |
| TUBB4B | BELANTAMAB MAFODOTIN | Approved |  | 0.036557282 |
| TUBB4B | VERUBULIN | Not Approved |  | 0.041431586 |
| TUBB4B | ANG1005 | Not Approved | antineoplastic agent | 0.036557282 |
| TUBB4B | RG-7600 | Not Approved |  | 0.038842112 |
| TUBB4B | INDIBULIN | Not Approved |  | 0.038842112 |
| TUBB4B | TUSAMITAMAB RAVTANSINE | Not Approved |  | 0.038842112 |
| TUBB4B | COMBRETASTATIN A4 | Not Approved |  | 0.276210574 |
| TUBB4B | BRENTUXIMAB VEDOTIN | Approved | antineoplastic agent | 0.034526322 |
| TUBB4B | MAYTANSINOL | Not Approved |  | 0.077684224 |
| TUBB4B | VINORELBINE | Approved | antineoplastic agent | 0.047805676 |
| TUBB4B | ADO-TRASTUZUMAB EMTANSINE | Approved | antineoplastic agent | 0.036557282 |
| TUBB4B | CURCUMIN | Approved |  | 0.018832539 |
| TUBB4B | PF-06263507 | Not Approved |  | 0.038842112 |
| TUBB4B | T-900607 | Not Approved |  | 0.041431586 |
| TUBB4B | PLINABULIN | Not Approved |  | 0.041431586 |
| TUBB4B | VINCRISTINE | Approved | antineoplastic agent | 0.030690064 |
| TUBB4B | VINFLUNINE | Approved |  | 0.041431586 |
| TUBB4B | PRALUZATAMAB RAVTANSINE | Not Approved |  | 0.038842112 |
| TUBB4B | VINBLASTINE | Approved | Antineoplastic Agents | 0.054835923 |
| TUBB4B | LAROTAXEL | Not Approved |  | 0.041431586 |
| TUBB4B | CROLIBULIN | Not Approved |  | 0.041431586 |
| TUBB4B | DAVUNETIDE | Not Approved |  | 0.041431586 |
| TUBB4B | PACLITAXEL POLIGLUMEX | Not Approved |  | 0.041431586 |
| TUBB4B | MEBENDAZOLE | Approved | antineoplastic agent | 0.124294758 |
| TUBB4B | MILATAXEL | Not Approved | antineoplastic agent | 0.036557282 |
| TUBB4B | APRUTUMAB IXADOTIN | Not Approved |  | 0.038842112 |
| TUBB4B | CYCLOSTREPTIN | Not Approved |  | 0.077684224 |
| TUBB4B | CANTUZUMAB MERTANSINE | Not Approved |  | 0.038842112 |
| TUBB4B | FOSBRETABULIN TROMETHAMINE | Not Approved |  | 0.041431586 |
| TUBB4B | GLEMBATUMUMAB VEDOTIN | Not Approved |  | 0.038842112 |
| TUBB4B | PATUPILONE | Not Approved |  | 0.041431586 |
| TUBB4B | POLATUZUMAB VEDOTIN | Approved |  | 0.032709147 |
| TUBB4B | MAYTANSINE | Not Approved |  | 0.077684224 |
| TUBB4B | KOS-1584 | Not Approved |  | 0.041431586 |
| TUBB4B | ANVATABART OPADOTIN | Not Approved |  | 0.038842112 |
| TUBB4B | PINATUZUMAB VEDOTIN | Not Approved |  | 0.038842112 |
| TUBB4B | PACLITAXEL | Approved | for treatment of peripheral arterial disease (PAD),DMARD,antiinflammatory agent,antineoplastic agent | 0.073446903 |
| TUBB4B | VINBLASTINE SULFATE | Approved |  | 0.03107369 |
| TUBB4B | EPOTHILONE D | Approved | antineoplastic agent | 0.038842112 |
| TUBB4B | LEXIBULIN | Not Approved |  | 0.041431586 |
| TUBB4B | LIFASTUZUMAB VEDOTIN | Not Approved |  | 0.038842112 |
| TUBB4B | LADIRATUZUMAB VEDOTIN | Not Approved |  | 0.038842112 |
| FOLR3 | PEMETREXED DISODIUM | Approved | Antineoplastic Agents,antineoplastic agent | 0.803135362 |
| FOLR3 | PETESICATIB | Not Approved | indication | 3.480253237 |
| FOLR3 | CITIBRASINE | Not Approved |  | 10.44075971 |
| FOLR3 | VBY-825 | Not Approved |  | 2.088151942 |
| FOLR3 | CITRUSININE II | Not Approved |  | 5.220379856 |
|  |  | regulatory approval |  | interaction score |
